# RPL22 is a tumor suppressor in MSI-high cancers and a key splicing regulator of MDM4

**DOI:** 10.1101/2023.12.10.570873

**Authors:** Hannah N.W. Weinstein, Kevin Hu, Lisa Fish, Yih-An Chen, Paul Allegakoen, Keliana S. F. Hui, Julia H. Pham, Maria B. Baco, Hanbing Song, Andrew O. Giacomelli, Francisca Vazquez, Mahmoud Ghandi, Hani Goodarzi, Franklin W. Huang

## Abstract

Microsatellite instability high (MSI-H) tumors are malignant tumors that, despite harboring a high mutational burden, often have intact *TP53*. One of the most frequent mutations in MSI-H tumors is a frameshift mutation in *RPL22*, a ribosomal protein. Here, we identified *RPL22* as a modulator of *MDM4* splicing through an alternative splicing switch in exon 6. *RPL22* loss increases *MDM4* exon 6 inclusion, cell proliferation, and augments resistance to the MDM inhibitor Nutlin-3a. RPL22 represses expression of its paralog, RPL22L1, by mediating the splicing of a cryptic exon corresponding to a truncated transcript. Therefore, damaging mutations in RPL22 drive oncogenic MDM4 induction and reveal a common splicing circuit in MSI-H tumors that may inform therapeutic targeting of the MDM4-p53 axis and oncogenic RPL22L1 induction.

## Introduction

Microsatellite instability high (MSI-H) tumors are a broad group of cancers that contain 30% or more mutations in microsatellites and are driven by somatic or germline mutations in DNA mismatch repair machinery^1,2^. There is a high MSI-H prevalence in uterine corpus endometrial carcinoma (28.3%), stomach adenocarcinoma (21.9%), and colon adenocarcinoma (16.6%)^3^. Despite a high tumor mutational burden, MSI-H tumors do not frequently harbor mutations in *TP53*, suggesting MSI-H tumors might harbor alternative mechanisms to inactivate p53^4^. Notably, MSI-H tumors have a frequent frameshift mutation (*p.K15fs*) in a microsatellite repeat site of the ribosomal protein *RPL22*^5–7^. *RPL22* and its paralog *RPL22L1* form part of the large subunit of the 60S ribosome and are known to affect protein synthesis as well as splicing of genes and transcription factors that affect development^8,9^ and tumorigenesis^10–13^. *RPL22* and *RPL22L1* expression levels are tightly linked with lower levels of one paralog resulting in higher expression of the other^8,14^. Of note, high RPL22L1 expression is specifically associated with increased *MDM4* exon 6 inclusion and *MDM4* dependency in Cancer Cell Line Encyclopedia (CCLE) data^5^. MDM4 protein expression is mediated through alternative mRNA splicing of its sixth exon, which leads to two major isoforms, the exon 6-inclusive *MDM4*-FL (full-length) and the exon 6-exclusive *MDM4*-S (short), which is unstable and prone to degradation through nonsense-mediated decay (NMD)^15–17^. The candidate regulators of this major splicing switch that control MDM4 activity have included SRSF3, ZMAT3, and PRMT5, among others^15,17,18^. However, to date, no mutational events in cancer have been associated with this alternative splicing event.

## Results

### Deleterious RPL22 alterations are common in MSI-H cancers and associated with specific splicing events of MDM4 and RPL22L1

We first considered the prevalence of *RPL22 p.K15fs* mutations in MSI-H tumors. We found 68% of MSI-H cell lines in the Cancer Cell Line Encyclopedia, as previously reported^5^, and 52% of MSI-H tumors in The Cancer Genome Atlas Program (TCGA) harbor this specific mutation (Supp. Data 1). The *RPL22 p.K15fs* mutation is also present at high rates in specific MSI-H cell lines and tumors, including colon (77%), endometrial (72%), gastric (67%), and ovarian (67%) cell lines in the CCLE, and colon adenocarcinoma (COAD, 48%), stomach adenocarcinoma (STAD, 80%) and uterine corpus endometrial carcinoma (UCEC, 59%) in the TCGA (Fig. 1a). Additionally, heterozygous *RPL22* copy number loss was common across many other tumor types, including kidney chromophobe (KICH, 79%), pheochromocytoma and paraganglioma (PCPG, 62%), adrenocortical carcinoma (ACC, 39%), and liver hepatocellular carcinoma (LIHC, 42%) (Fig. 1b). We found that *RPL22* frameshift mutations were mutually exclusive with deleterious *TP53* alterations (Supp. Fig. 1a-b).

**Figure 1.**
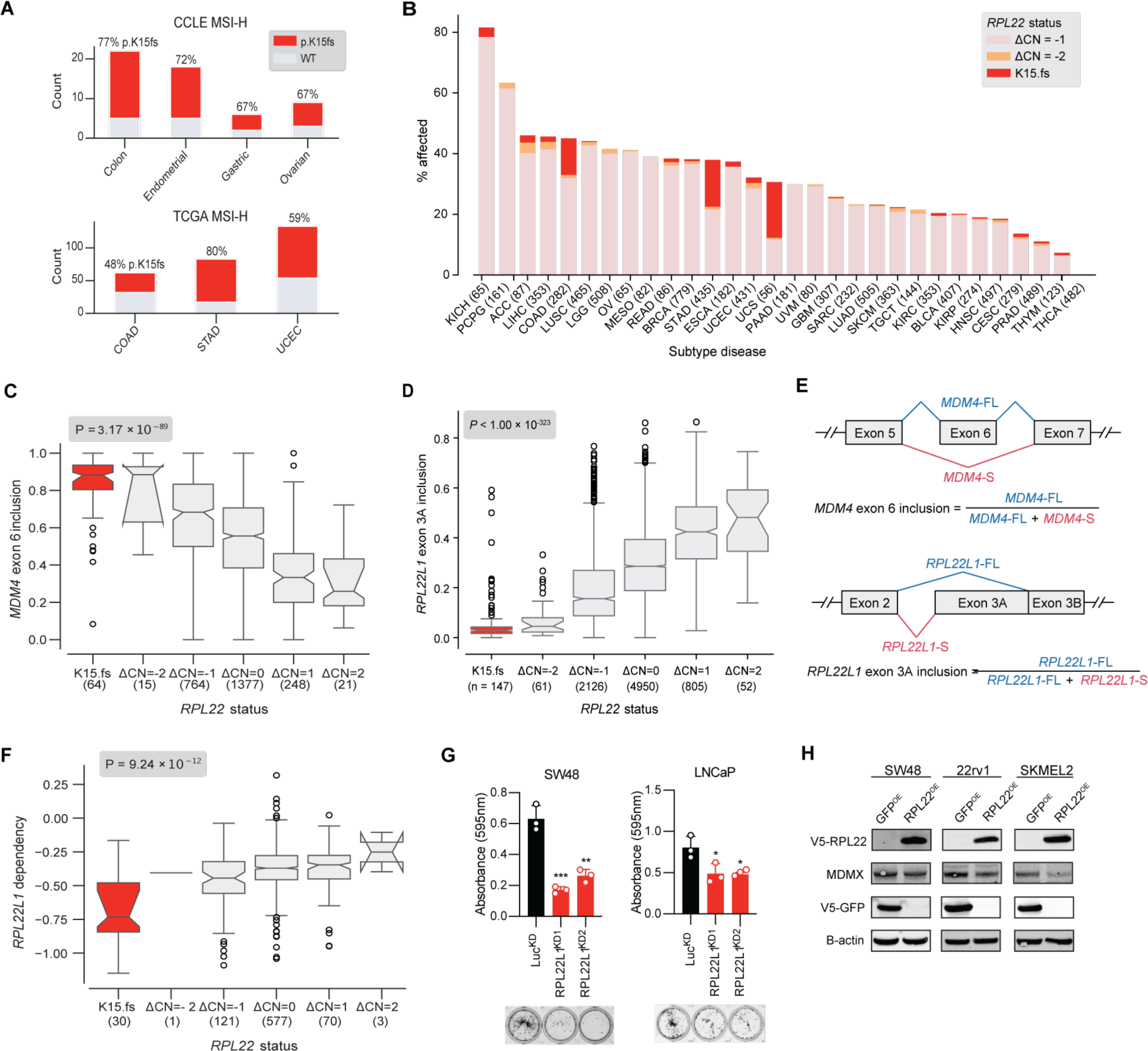
*RPL22* genomic alterations are common in cancer and associated with changes to specific transcripts of *MDM4* and *RPL22L1* mRNA. (**A**) Frequencies of *RPL22 p.K15fs* frameshift mutation in MSI-H cell lines in the CCLE and MSI-H tumors in the TCGA (COAD = colon adenocarcinoma; STAD = stomach adenocarcinoma; UCEC = uterine corpus endometrial carcinoma). (**B**) Proportion of *RPL22* copy number loss and truncating mutations across TCGA tissue types. (**C**) Association between decreased inclusion of *MDM4 exon 6* and deleterious alterations to *RPL22* in TCGA samples. (**D**) Association between decreased inclusion of *RPL22L1* exon 3A and deleterious alterations to *RPL22* in TCGA samples. (**E**) Schematic of splicing switches responsible for the major *MDM4* (top) and *RPL22L1* (bottom) isoforms. (**F**) Association between *RPL22L1 dependency* and deleterious alterations to *RPL22* in CCLE samples. (**G**) Focus formation assay of three microsatellite instable (MSI), *RPL22* mutant cell lines with shRNA knockdown of *RPL22L1*. Crystal violet staining was quantitated 10-12 days post seeding and compared to control by Student t test (p-value: SW48 RPL22L1^KD1^ = 0.0008; SW48 RPL22L1^KD2^ = 0.0025; LNCap RPL22L1^KD1^ = 0.0400; LNCap RPL22L1^KD2^ = 0.0155). Error bars represent the mean ± S.D of three replicates (* p < 0.05, ** p < 0.01, *** p < 0.001, **** p < 0.0001.) Data shown are representative of two independent experiments. (**H**) Western blot of MSI, RPL22 mutant cell lines with overexpression of RPL22 and knockdown of RPL22L1. Immunostained for V5-RPL22, MDM4, V5-GFP, and B-actin (30ug protein/lane).

As *RPL22* is recurrently mutated in MSI-H tumors, we tested for specific splicing events and gene expression changes related to *RPL22* loss. *RPL22* copy number loss and *RPL22* frameshift mutations were strongly correlated with *MDM4* exon 6 inclusion in TCGA samples (Fig. 1c). Deleterious alterations in *RPL22* were also associated with decreased inclusion of an alternative 3’-splicing event in *RPL22L1* adjacent to exon 3 (Fig. 1d). This transcript is predicted to result in a premature stop codon in exon 3 leading to a 63 amino acid protein compared to the full-length RPL22L1 isoform (122 amino acids). The exon affected by this alternative 3’-splicing event is defined here as exon 3A (Fig. 1e). Taken together these results suggest that *RPL22* loss is associated with *MDM4* exon 6 inclusion and regulates its paralog RPL22L1 protein expression through exon 3A alternative splicing.

### RPL22 and RPL22L1 exhibit paralog synthetic lethality

As paralogs, RPL22 and RPL22L1 share 73% amino acid identity^14^. *RPL22L1* has previously been identified as a paralog synthetic lethality in the context of *RPL22* mutation^5,19^. In the context of MSI-H tumors, *RPL22L1* is the second-highest scoring synthetic lethality after *WRN*, likely due to the high frequency of *RPL22* frameshift mutations in MSI-H tumors^20^. To confirm the *RPL22/RPL22L1* synthetic lethality relationship, synthetic lethality was tested in the updated CRISPR-Avana data set using 808 cancer cell lines. *RPL22L1* dependency was highly correlated with *RPL22* mutational status (Fig. 1g). Knockdown of *RPL22L1* using two independent shRNAs in two *RPL22* mutant MSI-H cell lines (Fig. 1h; Supp. Table 1) and *RPL22* knockout in one MSS cell line (Supp. Fig. 2a) validated the synthetic lethality. This synthetic lethality was not dependent on intact p53 as isogenic TP53-wild-type and TP53-null RKO cells harboring *RPL22fs* mutations were both sensitive to *RPL22L1* knockdown (Supp. Fig. 2b). Furthermore, overexpression of *RPL22* was sufficient to rescue the synthetic lethality (Supp. Fig. 2c).

### Modulation of RPL22 regulates a limited gene set, including RPL22L1

We next tested whether RPL22L1 regulates expression of RPL22. Overexpression of *RPL22L1* in an MSS cell line led to a robust increase in cell proliferation (Supp. Fig. 3a) and a 1.9-fold decrease in *RPL22* mRNA expression (Supp Fig. 3b), but knockdown of *RPL22L1* did not increase *RPL22* mRNA expression (Supp. Fig. 3b). However, overexpression of *RPL22* in a MSI-H cell line led to a significant 3.8-fold decrease in *RPL22L1* mRNA expression while CRISPR/Cas9 knockout of *RPL22* in two MSS cell lines resulted in a significant ∼10-fold increase in *RPL22L1* mRNA expression (Fig. 2a). These results suggest that RPL22 regulates *RPL22L1* mRNA expression, while modulation of *RPL22L1* does not strongly affect *RPL22* mRNA expression.

**Figure 2.**
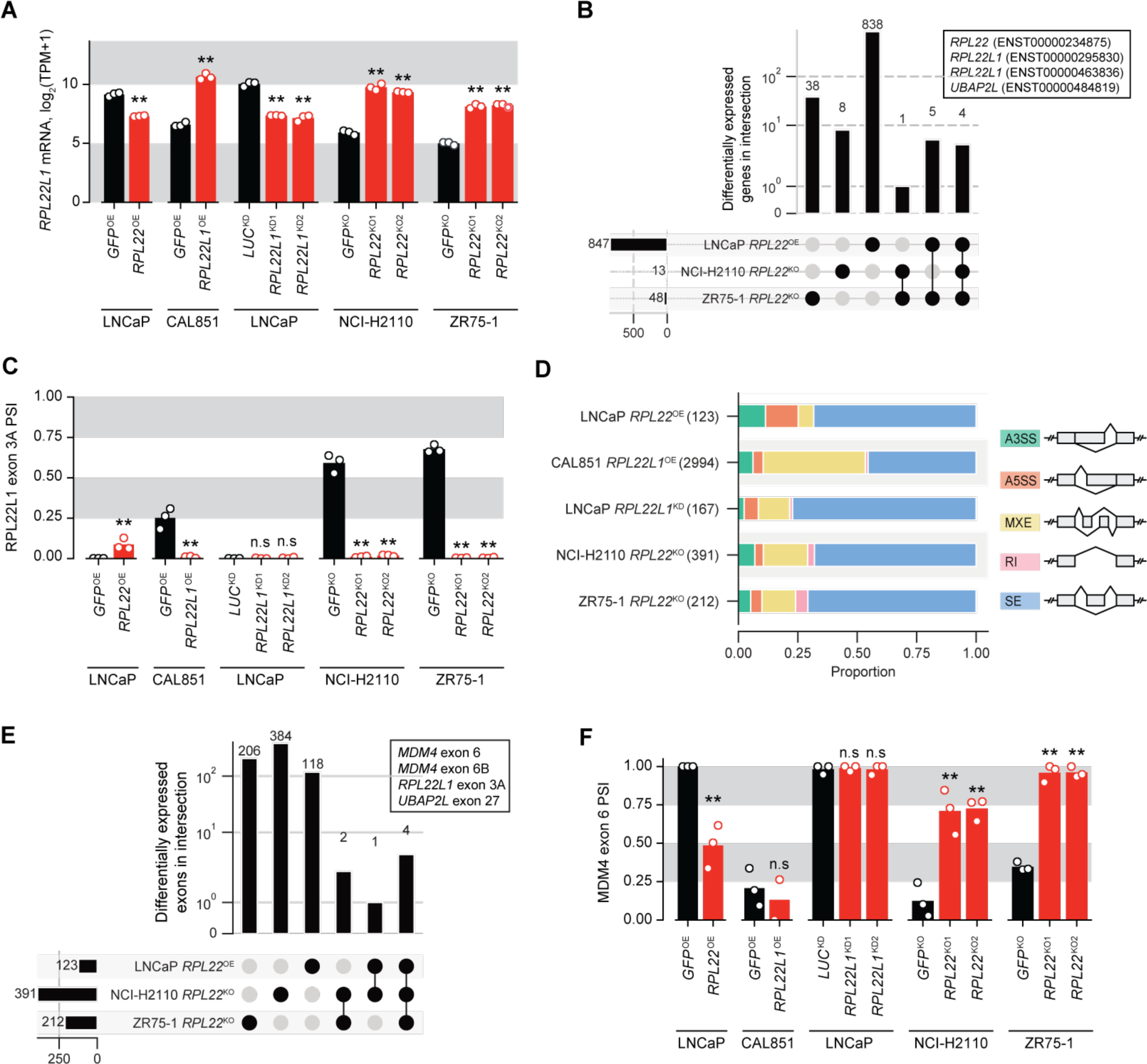
*MDM4* and *RPL22L1* are splicing targets of *RPL22*. (**A**) Total *RPL22L1* mRNA expression across experiments. (**B**) Common differentially expressed genes across *RPL22* knockout experiments. (**C**) Inclusion proportions of *RPL22L1* exon 3A across experiments. *RPL22* knockdown reduces *RPL22L1* exon 3a inclusion. (**D**) Changes in splicing modes across experiments as indicated by significant differentially spliced variants. (**E**) Common splicing changes across *RPL22* knockout experiments. (**F**) Inclusion levels of *MDM4* exon 6 across experiments. RPL22 overexpression reduces *MDM4* exon 6 inclusion and *RPL22* knockdown increases *MDM4* exon 6 inclusion; RPL22L1 overexpression and knockdown does not affect *MDM4* exon 6 inclusion.

**Figure 3.**
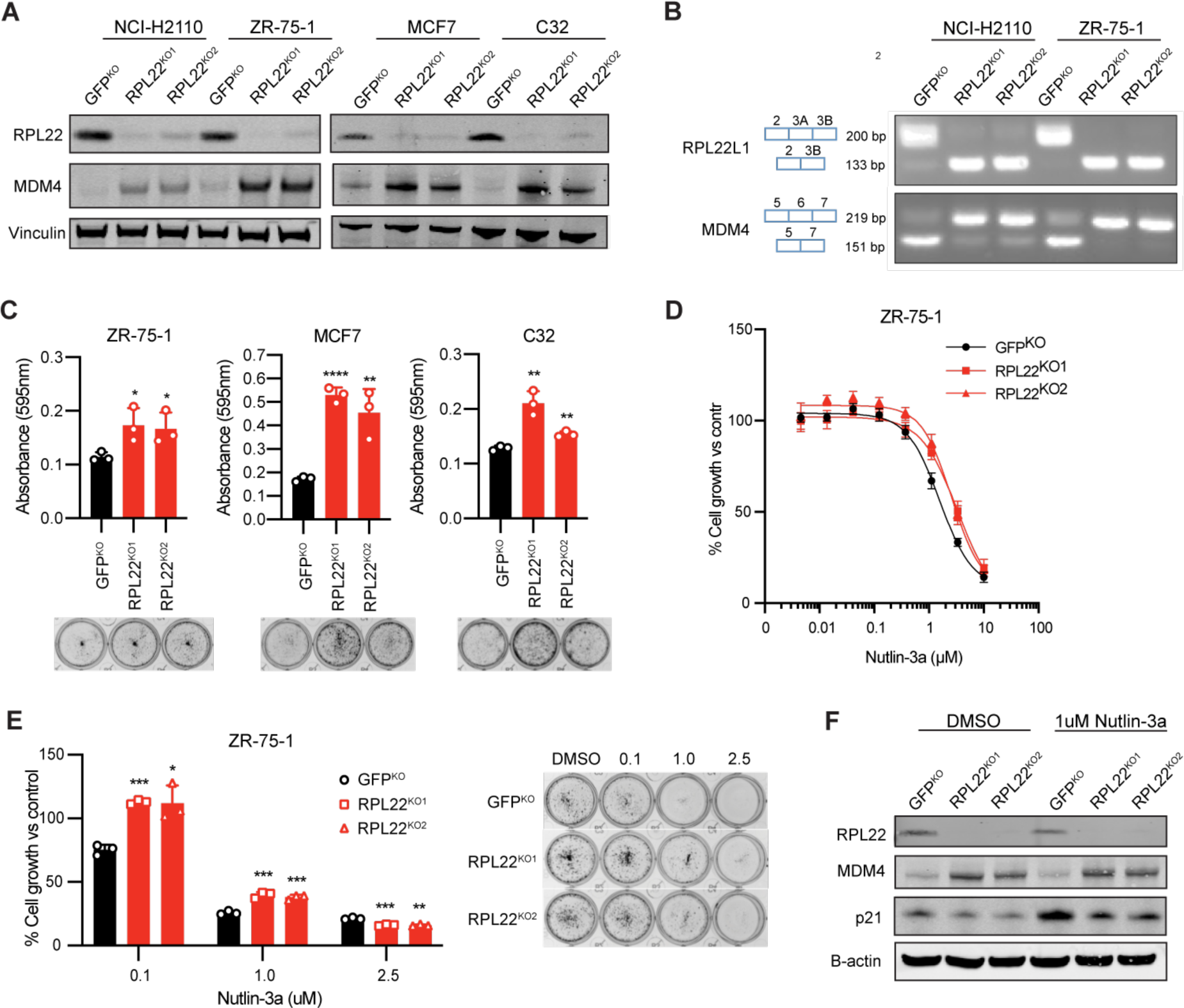
Loss of *RPL22* increases *MDM4* expression and augments resistance to Nutlin-3a. **(A)** Western blot analyses of cell lines with CRISPR-Cas9 knockout of *RPL22* with immunostaining for MDM4 (30ug protein/lane). (**B**) RT-PCR analysis of cell lines with CRISPR-Cas9 knockout of *RPL22.* (**C**) Focus formation assay of cell lines with CRISPR-Cas9 knockout of *RPL22*. Crystal violet staining was quantitated 7-14 days post seeding and compared to control by Student t test (p-value: ZR-75-1 RPL22^KO1^ = 0.0377; ZR-75-1 RPL22^KO2^ = 0.0464; MCF7 RPL22^KO1^ = <0.0001; MCF7 RPL22^KO2^ = 0.0089; C32 RPL22^KO1^ = 0.0033; C32 RPL22^KO2^ = 0.0029). Error bars represent the mean ± S.D of three replicates (* p < 0.05, ** p < 0.01, *** p < 0.001, **** p < 0.0001.) Data shown are representative of at least three independent experiments. (**D**) Nutlin-3a treatment of cell lines (ZR-75-1, 5000 cells/well) with CRISPR-cas9 knockout of *RPL22*. Luminescence was quantified 6 days post-seeding and compared to control by Student t test. Error bars represent the mean ± S.D of six replicates. Data shown are representative of three independent experiments. IC50 were as follows: GFP^KO^ = 1.566 uM, RPL22^KO1^ = 3.297 uM, RPL22^KO2^ = 2.574 uM. (**E**) Focus formation assay with nutlin-3a treatment of cell lines (ZR-75-1, 7.5k cells/well) with CRISPR-cas9 knockout of *RPL22*. Crystal violet staining was quantitated 14 days post-seeding and compared to control by Student t test (p-value: 0.1 uM nutlin-3a: ZR-75-1 RPL22^KO1^ = 0.0001; ZR-75-1 RPL22^KO2^ = 0.0126; 1.0 uM nutlin-3a: ZR-75-1 RPL22^KO1^ = 0.0008; ZR-75-1 RPL22^KO2^ = 0.0008; 2.5 uM nutlin-3a: ZR-75-1 RPL22^KO1^ = 0.0010; ZR-75-1 RPL22^KO2^ = 0.0015). Error bars represent the mean ± S.D of three replicates (* p < 0.05, ** p < 0.01, *** p < 0.001, **** p < 0.0001.) Data shown are representative figures of three independent experiments. (**F**) Western blot of cell line (ZR-75-1) with CRISPR-Cas9 knockout of *RPL22* with immunostaining for p21 after 1 week of 1uM nutlin-3a treatment (30ug protein/lane).

Given modulation of *RPL22* regulates *RPL22L1* mRNA expression, we performed intersection analyses across *RPL22* and *RPL22L1* modulation experiments in MSI-H and MSS cell lines to identify genes affected by RPL22 or RPL22L1 perturbation (Supp. Table 1). Across all *RPL22* overexpression and knockout experiments, four common differentially expressed transcripts were identified: *RPL22*, two protein coding transcripts of *RPL22L1*, and *UBAP2L* (Fig. 2b), indicating modulation of *RPL22* leads to a highly specific change in expression of its paralog, RPL22L1. UBAP2L (ubiquitin-associated protein 2-like) is an RNA-binding protein that promotes translation and was not further investigated here^21^. Perturbation of *RPL22L1* resulted in changes in over 400 genes that did not include *RPL22* (Supp Fig. 3c-d).

An intersection analysis between genes that exhibited differential expression and those with differentially spliced exons within each experiment revealed large, shared gene sets in the *RPL22L1* modulation experiments. Over half of the differentially spliced genes were also differentially expressed following *RPL22L1* overexpression (Supp Fig. 3e). However, smaller intersections were present among the *RPL22* experiments, with far greater numbers of differentially spliced genes, compared to those that were differentially expressed, associated with *RPL22* knockout (Supp Fig. 3e). Gene set analysis indicated that loss of *RPL22L1* led to upregulation of hallmark p53 genes, whereas overexpression of *RPL22L1* led to downregulation as well as converse effects on other pathways, such as chromosome condensation and segregation (Supp Fig. 3f). Overall, the results suggest that RPL22L1 modulation affects the expression and splicing of many genes involved in cell proliferation, whereas RPL22 directly regulates the expression of a more limited gene set that includes *RPL22L1*.

### RPL22 controls RPL22L1 expression via a cryptic alternative 3’ splice site and MDM4 expression through splicing of MDM4 exon 6

Based on our identification of a cryptic alternative 3’ splicing event (Supp Fig. 3g) resulting in exon 3A inclusion that is highly correlated with *RPL22* loss, we tested whether RPL22 modulates *RPL22L1* mRNA expression by regulating splicing of *RPL22L1* pre-mRNA. Expression of *RPL22* in an MSI-H *RPL22*-deficient (LNCaP) cells and knockout of *RPL22* in MSS *RPL22*-wildtype (NCI-H2110 and ZR75-1) cells significantly increased *RPL22L1* exon 3A inclusion, while *RPL22* knockout robustly decreased *RPL22L1* exon 3A inclusion (Fig. 3c).

Differentially expressed splicing events in the context of *RPL22* or *RPL22L1* modulation further defined the splicing events that occur in the setting of *RPL22* loss. Across all modulation experiments the most common splicing events were skipped exons with similar numbers of inclusion and exclusion events. In addition, overexpression of *RPL22L1* in MSS RPL22 intact cells (CAL851) coincided with altered splicing in large numbers of mutually exclusive exons (Fig. 2d). Over expression or knockout of *RPL22* led to specific differentially spliced exon events of *MDM4* exon 6, *MDM4* exon 6B, *RPL22L1* exon 3A, and *UBAP2L* exon 27 (Fig. 2e). Between the *RPL22L1* modulation experiments, 14 genes (SMPD4, FN1, EEF1D, DERL2, RPS24, ATXN2, H2AFY, FN1, ASPH, MYO5A, USMG5, RPS24, SCRIB, DERL2) were shared (Supp Fig. 3d). Modulation of *RPL22* had no significant impact on *MDM4* and *MDM2* mRNA levels (Supp. Fig. 3h-i), suggesting a primary role in modulating splicing. Overall, we conclude that RPL22 modulation leads to precise changes in the pre-mRNA splicing of *MDM4* exon 6, *RPL22L1* exon 3A, and *UBAP2L* exon 27.

RPL22L1 was not necessary or sufficient to promote *MDM4* exon 6 inclusion (Fig. 2f). However, overexpression of *RPL22* led to a marked decrease in *MDM4* exon 6 inclusion and *RPL22* knockout robustly increased *MDM4* exon 6 inclusion across multiple cell lines (Fig. 2f). Together, these data further suggest that RPL22 tightly regulates the alternative splicing and expression of both RPL22L1 and MDM4.

### RPL22 functions as a tumor suppressor through regulation of MDM4

Based on our RNAseq analysis showing an increase in *MDM4* exon 6 inclusion and *RPL22L1* exon 3A exclusion in the context of *RPL22* loss, we hypothesized MDM4 and RPL22L1 protein levels increase when *RPL22* is lost. In CCLE data, MDM4 protein was most strongly correlated with both *MDM4* mRNA and *MDM4* exon 6 inclusion (Supp. Fig. 4a), suggesting MDM4 protein expression occurs in the context of MDM4-FL transcripts as previous studies have shown^15^. To test the loss of RPL22, we performed CRISPR/Cas9 knockout of *RPL22* in four *RPL22* wildtype MSS cell lines (ZR-75-1, NCI-H2110, C32, and MCF7), and observed a marked decrease in RPL22 protein expression and robust increase in MDM4 protein expression (Fig. 3a). The two specific splicing events of *MDM4* exon 6 inclusion and the alternative 3’ splicing event in *RPL22L1* identified through RNAseq were validated using reverse-transcriptase polymerase chain reaction (RT-PCR) analysis following CRISPR/Cas9 knockout of *RPL22* (Fig. 3b). Furthermore, colony formation assays showed a significant increase in cell proliferation in the context of *RPL22* loss compared to control in p53 wildtype cells (Fig. 3c) but not in p53 mutant cells (Supp. Fig. 4b).

Since MDM4 is a well-known inhibitor of p53, we hypothesized the increase in cell growth in the context of *RPL22* loss was driven primarily by increased MDM4 protein levels. To test whether increased MDM4 levels contributed to increased cell growth in the context of *RPL22* loss, cells were treated with Nutlin-3a, an MDM2 inhibitor with some activity against MDM4^22^. Nutlin-3a sensitivity is highly correlated with the full length version of MDM4 (MDM4-FL) and is the top drug sensitivity correlated with *MDM4* exon 6 inclusion among a large panel of drugs^5^. *TP53* wildtype MSS cells with CRISPR/Cas9 knockout of *RPL22* showed resistance to Nutlin-3a in short and long-term growth assays (Fig. 3d-e). The results suggest that while MDM4-FL levels appear to be a biomarker for Nutlin-3a sensitivity, loss of RPL22 and overexpression of MDM4 promotes resistance to Nutlin-3a. No difference in cell growth was observed in p53 mutant cells treated with Nutlin-3a (Supp. Fig. 4c-d). While induction of p53 protein expression was not observed in the context of one week of Nutlin-3a treatment and *RPL22* knockout (Supp. Fig. 4e), protein expression of p21, a downstream target of p53, was downregulated (Fig. 3f), consistent with p53 inhibition through MDM4. Quantitative polymerase chain reaction (qRT-PCR) of downstream targets of p53, including BBC3, p21 and MDM2, showed significant induction following Nutlin-3a treatment with less induction in the context of *RPL22* loss but no significant difference was consistently observed after 24 hours or one week of treatment (Supp. Fig. 4f).

Since RPL22 and RPL22L1 are known to regulate pre-mRNA splicing by binding to the mRNA of their targets^8^, and experiments suggested that RPL22 was the paralog responsible for modulating the splicing of *MDM4*, *RPL22L1*, and *UBAP2L*, the predicted hairpin structures in *MDM4* proximal to exon 6A and in *RPL22L1* proximal to exon 3A, where RPL22 might bind, were of particular interest (Supp. Fig. 5-6). We performed CLIP-seq on *TP53* wildtype MSS cells to characterize the genome-wide binding specificity of RPL22 and identified significant peaks in the 3’ UTR of *MDM4* and *UBAP2L* but no such peaks in *RPL22L1* (Fig. 4a). We also confirmed binding of RPL22 to *MDM2* mRNA as previous studies have shown (Supp. Fig. 7)^23^. Region enrichment analysis uncovered a binding preference for the transcription termination site (Fig. 4a-b), and gene set enrichment analysis of genes containing peaks uncovered strong enrichment (p<1*10^-7^) for genes involved in histone modification, regulation of transcription, and RNA splicing (Fig. 4c). Intersections between peak-containing genes and those exhibiting significant differential splicing across previous *RPL22*- and *RPL22L1*-modulating experiments also revealed sizable overlaps (Fig. 4d). Taken together, these results indicate that RPL22 modulates the expression of *MDM4* and *RPL22L1* through splicing, ultimately promoting cell proliferation and survival (Fig. 4e).

**Figure 4.**
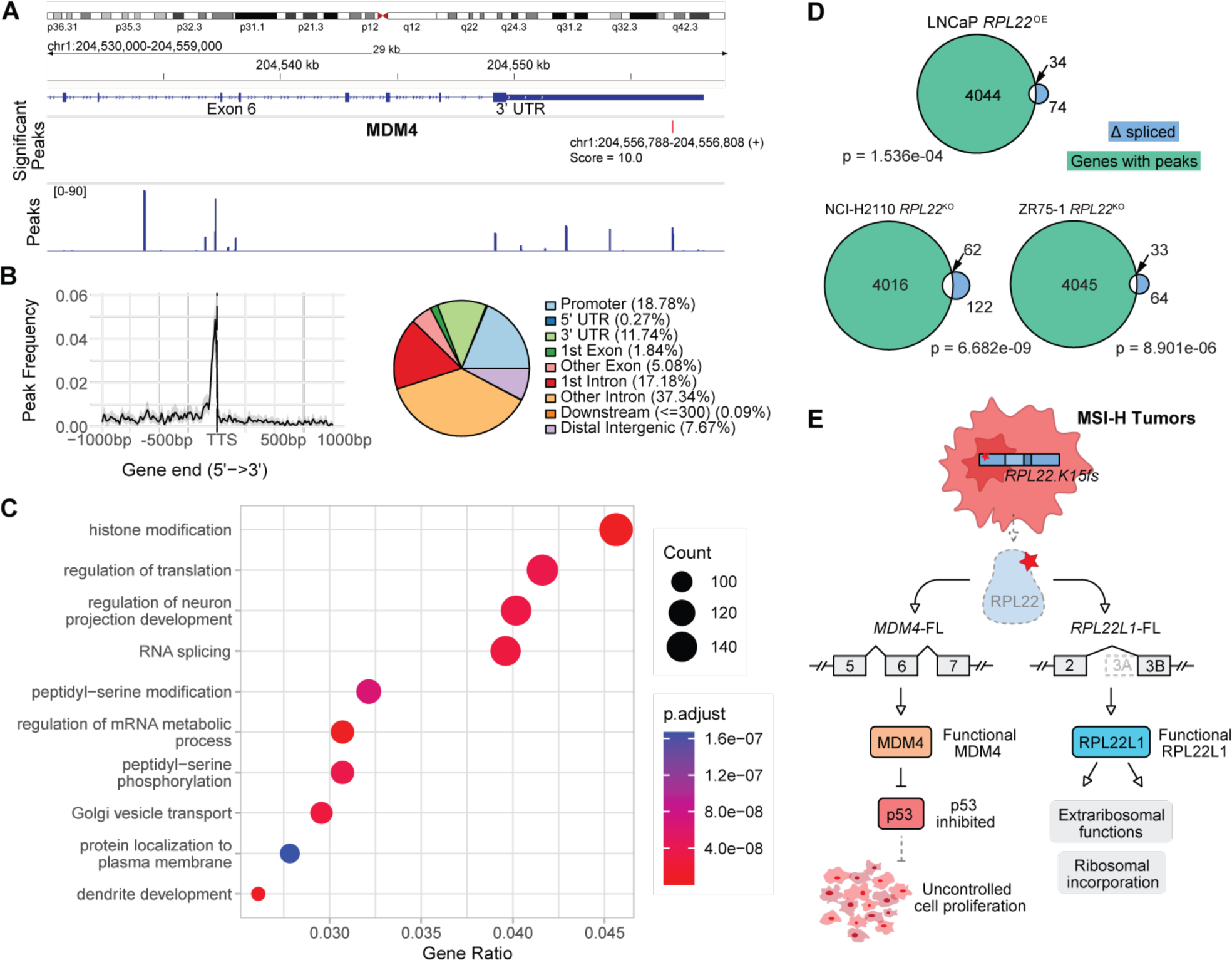
Characterization of RPL22 binding preference with CLIP-seq. (**A**) Binding of RPL22 to the 3’ UTR and exon 6 of *MDM4* as determined by peak calling on CLIP-seq measurements from ZR-75-1 cells. (**B**) Left: Peak frequencies relative to the transcription termination site (TTS). Right: Breakdown of peak-containing regions. (**C**) Gene set enrichment analysis of RPL22-bound peaks. (**D**) Overlaps between genes containing RPL22-bound peaks and differentially spliced genes across *RPL22* modulation experiments. Total number of genes and hypergeometric test for each cell line: LNCaP: 15602 genes, p = 1.536e-04; NCI-H2110: 15226 genes, p = 6.682e-09; ZR75-1: 15690 genes, p = 8.901e-06. (**E**) Schematic model of loss of *RPL22* frequent in MSI-H tumors that results in a splicing-mediated switch to the functional isoforms of *MDM4* and *RPL22L1*. The functional isoform of *MDM4* inhibits p53, and increased levels of RPL22L1 compensate for RPL22 reduction in the ribosome and perform additional extra ribosomal functions.

## Discussion

In this study, we identify the activation of a common splicing switch through the loss of RPL22 and reveal it as a mechanism of MDM4 induction. *RPL22* loss increases inclusion of MDM4 exon 6, leading to a functional protein that is known to interact with MDM2 and together form a complex with p53 to inactivate and target p53 for degradation. We identify a physical interaction between RPL22 and the *MDM4* 3’-UTR, which may indicate additional regulatory mechanisms. Through these interactions, RPL22 may function to maintain the expression of the MDM4 short isoform (MDM4-S) in the balance between MDM4-FL/MDM4-S isoforms, thus providing a key mechanism to suppress MDM4-FL expression and an explanation for the lack of MDM4 protein expression in most somatic tissues^24^. Additionally, the *MDM4* 3’-UTR is a known site of microRNA regulation and singular nuclear polymorphisms (SNPs) that are associated with increased risk for multiple cancers^24^. It remains unclear whether RPL22 binds MDM4 alone or in conjunction with other primary splicing regulators. RPL22 is also thought to mediate cell proliferation through inhibitory RPL22 N-terminus binding to MDM2 but not through splicing regulation as we saw here^23^. In patients with MSI-H tumors, such as colon and stomach adenocarcinomas, which contain *RPL22fs* mutations in more than 70% of MSI-H cell lines and tumors in the CCLE and TCGA, our study suggests that tumorigenesis may in part be driven by *RPL22* mutation leading to MDM4 inhibition of p53.

We also show that *RPL22* loss leads to a splicing switch of its paralog RPL22L1, thereby highlighting how one ribosomal paralog regulates another. Inclusion of *RPL22L1* exon 3A is predicted to encode a non-functional protein lacking the RNA-binding domain as well as an mRNA transcript that is potentially subject to NMD^25^. The alternate 3’ splice site in the third exon of *RPL22L1* has also recently been described in the induction of glioblastoma cells under alternate regulation by serine/arginine-rich splicing factor 4 (SRSF4)^26^. Recent data indicate that this differentially spliced RPL22L1 has activity in glioblastoma cells and highlights another mechanism by which this transcript affects splicing^26^. Activation of RPL22L1 is also known to drive cell proliferation in ovarian tumors through the epithelial-to-mesenchymal transition^12^, indicating the importance of RPL22L1 activation through splicing in tumor progression. Although *RPL22* and *RPL22L1* share highly similar sequences and are functionally redundant in the large ribosome for ribosome biogenesis and protein synthesis^14^, these results suggest a wider range of nuclear effects of RPL22 and RPL22L1 on mRNA expression through splicing. Although we were unable to detect direct binding of RPL22 to RPL22L1, our findings combined with previous evidence of splicing-mediated autoregulation between *RPL22* paralogs in yeast^25,27–29^, and evidence of RPL22 repression of its paralog^14^, suggest RPL22 directly regulates its own paralog through this alternative 3’ splicing switch.

Given the high prevalence of *RPL22* frameshift mutations in MSI-high cancers, our results identify *RPL22* as a tumor suppressor gene that acts as a regulator of the p53 inhibitor MDM4 through modulation of its splicing. Our work suggests ribosomal genes that also function as mediators of splicing may serve as therapeutic splicing targets^30–34^. The identification of RPL22 as a key regulator of MDM4 function provides an additional pathway for development of MDM4-inhibiting treatments and adds the paralogs RPL22/RPL22L1 to other ribosomal proteins that modulate p53 activity in cancer.

## Acknowledgements

We thank all members of the Huang lab for their feedback and input.

## Author contributions

Conceptualization: HNW, KH, MG, FWH

Methodology: HNW, KH, YC, PA, KSH, JHP, LF, MB, HS, AG, FV, MG, HG, FWH

Software: KH, MG

Validation: HNW, KH, PA, YC, KSH, JHP

Formal analysis: HNW, KH, PA

Investigation: HNW, KH, PA, LF, MB, YC, KSH, JHP

Resources: HG, FWH

Data Curation: HNW, KH

Writing – original draft: HNW, FWH, KH

Writing – review & editing: HNW, KH, MG, FWH

Visualization: HNW, KH, FWH

Supervision: FWH

Project administration: FWH

Funding acquisition: FWH

## Data and materials availability

Raw RNAseq and CLIP-seq reads will be deposited in SRA under accession number [INSERT HERE]. All other data are available in the main text or the supplementary materials.

## Methods and Methods

### CCLE annotations

We sourced mutations and copy number estimates from the DepMap download portal (https://depmap.org/portal/download/) under the public 19Q4 release (CCLE_mutations.csv and CCLE_gene_cn.csv, respectively). We used the mutation classifications detailed in the Variant_annotation column. We also downloaded processed RNAseq estimates in the form of gene expression, transcript expression, and exon inclusion estimates from the DepMap data portal under the latest CCLE release (CCLE_RNAseq_rsem_genes_tpm_20180929.txt.gz, CCLE_RNAseq_rsem_transcripts_tpm_20180929.txt.gz, and CCLE_RNAseq_ExonUsageRatio_20180929.gct.gz, respectively). We also downloaded RPPA estimates (CCLE_RPPA_20181003.csv) and global chromatin profiling results (CCLE_GlobalChromatinProfiling_20181130.csv). Before performing subsequent analyses, we transformed transcript and gene expression TPMs by taking a log2-transform with a pseudocount of +1. Harmonized CCLE sample information and annotations are available in Supplementary Table 2.

### TCGA annotations

Pan-cancer TCGA sequencing data were sourced from the UCSC Xena browser (https://xenabrowser.net/datapages/?cohort=TCGA%20Pan-Cancer%20(PANCAN)&removeHub=https%3A%2F%2Fxena.treehouse.gi.ucsc.edu%3A443) as well as the Genomic Data Commons (https://gdc.cancer.gov/about-data/publications/PanCanAtlas-Splicing-2018). TCGA STAD (Stomach Adenocarcinoma) RNA-seq expression and somatic mutation data were sourced from cBioPortal (https://www.cbioportal.org/study/summary?id=stad_tcga). TCGA UCEC (Uterine Corpus Endometrial Carcinoma) RNA-seq expression and somatic mutation data were sourced from cBioPortal (https://www.cbioportal.org/study/summary?id=ucec_tcga). TCGA MSI annotations were sourced from Supplementary File 2 of Bonneville et al. 2017^35^. Harmonized TCGA sample information and annotations are available in Supplementary Table 3.

### Cell culture

Cells from various cancer types – breast, lung, prostate, colon, and skin – were cultured as monolayers. Of the included cell lines, SW48, 22rv1, SKMEL2, and LNCaP are known microsatellite instable (MSI) cell lines, while ZR-75-1, NCI-H2110, C32, MCF7, CAL851, and PC3 cells are microsatellite-stable (Table S1). RICR and RPCR cells were also created in a MSI background. ZR-75-1, C32, and MCF7 cells are *RPL22* and *TP53* wild-type, while NCI-H2110 cells are *RPL22* wild-type and contain a *TP53* p.R158P missense mutation. LNCaP cells contain a *RPL22 p.K15fs* frameshift mutation and are wild-type *TP53*. CAL851 cells are wild-type *RPL22* and contain a *TP53* p.K132E mutation. SW48, SKMEL2, and 22rv1 cells contain a *RPL22 p.K15fs* frameshift mutation. PC3 cells are TP53 null. All the cell lines listed above were purchased from the American Type Culture Collection (ATCC). ZR-75-1, NCI-H2110, C32, PC3 and LNCaP cells were cultured in RPMI (Gibco) with 10% FBS. SW48, SW1573, 22rv1, and CAL851 cells were cultured in DMEM (Gibco) with 10% FBS (Thermo Fisher). RICR (*TP53* mutant), RPCR (*TP53* wildtype), and MCF7 cells were cultured in EMEM (Gibco) with 10% FBS. RICR and RPCR were generated by transiently transfecting the parental *TP53* wildtype MSI RKO cells with sgRNAs (in pXPR003) and Cas9 (in pLX311), and then selecting for ∼2 weeks in 5 uM Nutlin-3. Thereafter, the cells were maintained in the absence of Nutlin-3 (DMEM + 10% FBS + Penn-Strep + Glutamine). The parental and p53-knockout cells were subsequently stably infected with Cas9 again in pLX311 and then with p53-WT or p53-MUT-P278A, making them all resistant to Blasticidin and Hygromycin. All cells were incubated at 37 °C with 5% CO2 and passaged twice a week using Trypsin-EDTA (0.25%) (Thermo Fisher). Cell identity was verified through the ATCC authentication service using short tandem repeat profiling and tested for mycoplasma using the Mycoalert Detection Kit (Lonza, LT07-118).

### Lentivirus Transduction

Lentiviruses were produced in 293T cells for all RNAi-induced knockdown, over-expression, and CRISPR-cas9 knockout experiments. To express shRNAs, cells were seeded at 4-5 x 10^5^ cells per well in a 6-well and subsequently transduced with lentivirus. The following shRNAs were obtained from the Genetic Perturbation Platform at the Broad Institute: plx-304 *lacZ* (LacZ^OE^) TCGTATTACAAGTCGTGACT; plx-304 *GFP* (GFP^OE^) ccsbBroad304_99986; plx-304 *RPL22* (RPL22^OE^) NM_000983; *shluc* (Luc^KD^) TRCN0000072243, CTTCGAAATGTCCGTTCGGTT; pLKO *RPL22L1* (RPL22L1^KD1^) TRCN0000247704: GAGGTCAACCTGGAGGTTTAA; pLKO *RPL22L1* (RPL22L1^KD2^) TRCN0000247705: CATTGAACGCTTCAAGAATAA. Puromycin (2 µg/mL) or blastocydin (5ug/ml) were used to select cells following incubation. shRNAs were obtained from the Genomics Perturbation Platform. GFP overexpression was verified through observation on a EVOS FL (Invitrogen) and *RPL22L1* knockdown was confirmed via western blot.

*RPL22* was targeted using single-guide RNA sequences inserted into the lentiCRISPR v2 plasmid purchased from Addgene and originally a gift from Feng Zhang (Addgene plasmid # 52961; http://n2t.net/addgene:52961; RRID:Addgene_52961)^36^. Six *RPL22* lentiCRISPR v2 constructs were tested and two, RPL22-1A-1 and RPL22-4A-1, were used based on western blot analysis. RPL22-1A-1 (RPL22^KO1^): CACCGCGCGGAGCCATACTAACCAC; RPL22-4A-1 (RPL22^KO2^): CACCGCATACTAACCACAGGAGCCA; control guide EGFP_sg6 (GFP^KO1^): GGTGAACCGCATCGAGCTGA^37^. Transduced cells were selected with puromycin (2 µg/mL) and western blot analysis was used to confirm *RPL22* knockout.

### Cell Proliferation Assays

Lentiviral transduced cells were plated for growth curve, drug curve and focus formation analysis. For growth and drug curves, cells were seeded at a density of 2-10 x 10^3^ cells in 96-well plates in RPMI (Gibco) with 10% FBS. Nutlin-3a (Tocris, 3984) drug dilutions were added after 24 hours and cells were allowed to grow for a subsequent 96 hours or stopped every 24 hours depending on the experiment. CellTiter-Glo was added to wells to stop the assay and luminescence recorded according to the manufacturer’s protocol on a SpectraMax (Molecular Devices). For focus formation analyses, cells were seeded in triplicate in 12-well plates at a density of 2-7.5 x 10^3^ in the appropriate media. If applicable, nutlin-3a was added the following day and replaced after seven days. Cells were fixed with 4% formaldehyde and stained with 0.5% crystal violet solution after 7-14 days depending on the cell line. For protein analysis and qPCR, cells were seeded in 6-well plates, treated with nutlin-3a after 24 hours with media replacement every three days, and collected following seven days of growth. Plates were imaged using a ChemiDoc Touch Gel Imaging System (Biorad) and subsequently de-stained using 10% acetic acid. Absorbance was measured at 595 nm using a SpectraMax (Molecular Devices). For all assays, statistical significance was calculated using unpaired student’s t-tests on GraphPad Prism 9 (GraphPad Software). Error bars represent standard deviation (s.d).

### RNA Sequencing, Reverse Transcription PCR, and Quantitative Real-time PCR

Total RNA was isolated using an RNAeasy Kit (Qiagen) according to the manufacturer’s instructions. For RNA sequencing at least 10,000 ng total RNA dissolved in ddH_2_O was extracted and sent to NovoGene. Libraries were prepared externally by NovoGene, sequenced on a NovaSeq 6000 PE150 and FASTQ files were provided.

For reverse transcription PCR (RT-PCR), 1 ug RNA was reverse transcribed with SuperScriptTM III First-Strand Synthesis System (ThermoFisher). PCRs were run using Q5® High-Fidelity DNA Polymerase. For RPL22L1 alternative splicing was visualized with the forward primer binding in exon 2, and the reverse primer binding in exon 3B. For MDM4, alternative splicing was visualized with forward primer binding in exon 5, and the reverse primer binding in exon 7. PCR was performed for 35 cycles (10s at 98℃; 20s at 64℃; 30s at 72℃) using a 100 ng cDNA template. Products were visualized on a 2% agarose gel. Products were visualized on a 2% agarose gel.

For quantitative real-time PCR (qPCR), 1 ug RNA was reverse transcribed with SuperScriptTM III First-Strand Synthesis System (ThermoFisher). Primers were purchased from Integrated DNA Technologies and sequences for all primers used in this study are included in Supplementary Table 2. To detect mRNA amplification reactions, qPCR was performed with the SYBR Green PCR master mix (Applied Biosystems) using a StepOnePlus Real-Time PCR system (Applied Biosystems). The ΔΔCt method was used to normalize the mRNA expression levels with TATA-box binding protein (TBP) as the house keeping gene.

### Western Blot Analysis

Cells were lysed in RIPA buffer with protease inhibitors (Roche, 4719956001), separated by SDS-PAGE and transferred to nitrocellulose membranes using the iBlot dry transfer system (Life Technologies). Membranes were cut, blocked with Intercept blocking buffers (LI-COR) and probed for antibodies against RPL22 (1:100, Santa Cruz, sc-136413), RPL22L1 (1:100, 1:5000, MyBioSource, MBS7050452), HdmX/MDMX (1:1000, Bethyl, A300-287A-M), p53 (1:200, Santa Cruz, sc-126), p53 (1:1000, Cell Signaling, 2524S), p21 (1:2000, Cell Signaling, 2946S), UBAP2L/NICE-4 (1:1000, Thomas Scientific, A300-534A-T) or Vinculin (1:500, Sigma, V9131). Secondary antibodies included goat anti-rabbit IRDye 800CW (LI-COR, 925- 32211), goat anti-mouse IRDye 800CW (LI-COR, 925-32210), and donkey anti-mouse IRDye 680 (LI-COR, 926-68072). Blots were imaged with an Odyssey Infrared Imager. All Western blots were repeated with separate biologically independent replicates.

### Transcript expression quantification

We used kallisto for transcript expression quantification. GRCh37 cDNA sequences for constructing the Kallisto indices were downloaded from the Ensembl FTP archives at ftp://ftp.ensembl.org/pub/release-75/fasta/homo_sapiens/cdna/Homo_sapiens.GRCh37.75.cdna.all.fa.gz. To construct the Kallisto index, place the cDNA FASTA file as-is into/data/raw/kallisto_homo_sapiens, and execute 1_kallisto_index.sh from the/scripts directory. Next, to quantify transcript expression, run 2_kallisto_quant.sh from/scripts/, which outputs estimates to/data/raw/kallisto_quant/. Kallisto fast enough to process each paired-end run in about half an hour on most modern machines. We ran Kallisto with arguments --bias -b 100 -t 6. Kallisto outputs raw transcript quantifications and bootstrap samples for use with Sleuth in differential expression analyses. To run Sleuth, see /notebooks/p1_kallisto-sleuth.ipynb. This was used to run differential expression analyses on each of the experiments, as well as annotate transcripts with matching genes and biotypes for downstream analyses.

### Signature gene set development and correlation analysis

Gene signatures were created for the four phenotypes (RPL22 KO, RPL22L1 KO, RPL22 overexpression, and RPL22L1 overexpression) in this study. Genes with log2 fold change > 1 and FDR q < 0.05 (compared to control samples, Wilcoxon rank-sum test) were included for each signature gene set. The ssGSEA module on GenePattern was then used to test the correlation of the gene signatures on the TCGA-STAD and TCGA-UCEC datasets with curated *TP53* and *RPL22* mutational status. Then the ssGSEA results .gct files were parsed through customized code to generate heatmaps and conduct correlation and statistical analysis.

### Splicing Quantification

To compute splicing levels and perform differential splicing analyses, we aligned FASTQ files with STAR, after which we used rMATS for splicing quantification. STAR alignment included the following process: First, we downloaded the Broad hg19 FASTA reference from console.cloud.google.com/storage/browser/_details/broad-references/hg19/v0/Homo_sapiens_assembly19.fasta, and the associated index from console.cloud.google.com/storage/browser/_details/broad-references/hg19/v0/Homo_sapiens_assembly19.fasta.fai. We also downloaded the Ensembl GTF file from ftp://ftp.ensembl.org/pub/release-75/fasta/homo_sapiens/dna/Homo_sapiens.GRCh37.75.dna.primary_assembly.fa.gz. STAR is a memory-heavy program, and building the index and alignments both require about 32-48 GB of RAM. An example script used for index construction on the Broad computing clusters is provided in /scripts/3_STAR_index.sh. Once the index was built, we ran two-pass alignments on the Broad computing clusters using 4_STAR_2pass.sh. Note the use of /scripts/fastq_samples.txt. We used the rMATS Docker image described at http://rnaseq-mats.sourceforge.net/rmatsdockerbeta. To run rMATS, we changed the DATA_PATH variable in /scripts/5_run_rmats.sh to the absolute path of the /data folder and executed the script.

### CLIP-seq

ZR-75-1 cells were crosslinked with 400mJ/cm2 254nm UV. Crosslinked cells were lysed with ice-cold lysis buffer (1X PBS, 0.1% SDS, 0.5% sodium deoxycholate, 0.5% IGEPAL CA-630) supplemented with 1x halt protease inhibitors (Pierce) and SuperaseIN (Invitrogen). After lysis, samples were treated with DNase I (Promega) at 37degC, 1000rpm 5 minutes. Each lysate sample was then split equally, and each portion treated with either a medium or low dilutions of a mix of RNase A and RNase I (Thermo; 3.3ng/ul RNase A and 1units/ul RNase I, and 0.67 RNase A and 0.2units/ul RNase I, respectively) at 37degC for 5 minutes. Lysate was immediately cleared by centrifuging at 21,000xg 4degC 20 minutes. The clarified lysate was added to protein G dynabeads (Invitrogen) that were pre-conjugated to an anti-RPL22 antibody (Santa Cruz sc-136413). Immunoprecipitation was performed at 4degC with end-over-end rotation for 1.5 hours. Beads were then washed 2X with ice-cold high salt wash buffer (5X PBS, 0.1% SDS, 0.5% sodium deoxycholate, 0.5% IGEPAL CA-630), 2X with ice-cold lysis buffer (1X PBS, 0.1% SDS, 0.5% sodium deoxycholate, 0.5% IGEPAL CA-630), and 2X with ice-cold PNK buffer (10mM Tris-HCl pH 7.5, 0.5% IGEPAL CA-630, 10mM MgCl2). The RNA was then end-repaired, and poly(A) tailed on-bead by treatment with T4 PNK (NEB) and then treatment with yeast poly(A) polymerase (Jena Bioscience). The RNA was 3’-end-labeled with azide-dUTP (TriLink biotechnologies) using yeast poly(A) polymerase (Jena Bioscience). The protein-RNA complexes were then labelled with IRDye800-DBCO (LiCor). The protein-RNA complexes were then eluted from beads in 1XLDS sample buffer (Invitrogen), resolved by running on a 4-12% Bis-Tris NuPAGE gel (Invitrogen), transferred to nitrocellulose, and imaged using an Odyssey Fc instrument (LiCor). The regions of interest were excised from the membrane and the RNA was isolated by Proteinase K (Invitrogen) digestion and subsequent pulldown with oligodT dynabeads (Invitrogen). After eluting from the oligodT beads, the RNA was immediately used for library preparation using the SMARTer smRNA-Seq Kit (Takara), with the following modifications. The poly(A) tailing step was omitted, and reverse transcription was performed with a custom RT primer. The PCR step was performed with indexed forward (i5) primers and a universal reverse (i7) primer. The libraries were PAGE purified and then sequenced on an Illumina HiSeq4000 instrument at the UCSF Center for Advanced Technologies.

### CLIP-seq Analysis

The raw CLIP-seq data were parsed via FastQC for quality control. Low-quality sequences, artifacts sequences, contaminated sequences and low-quality reads were filtered out via Clip tool kit^38^. Low-quality sequences before the adapters and universal adapter sequences were removed by Cutadapt trimmer. Reads with sufficient quality were then aligned to the human reference genome (hg38). The aligned sam file for each biological replicate was parsed and converted to .bed file using parseAlignment.pl, selectRow.pl and bed2rgb.pl. Using the combined results .bed files, peak calling was performed using Clip tool kit tag2peak.pl and statistical significance was set as p-value = 0.05. Integrative Genomics Viewer (IGV) was used to visualize the peaks^39^. Motif detection was conducted on the called peaks using HOMER^40^. Genomic annotations were obtained by extracting the coordinates of the significant peaks using bedtools, followed by gene ontology analysis.

---

**Figure S1.**
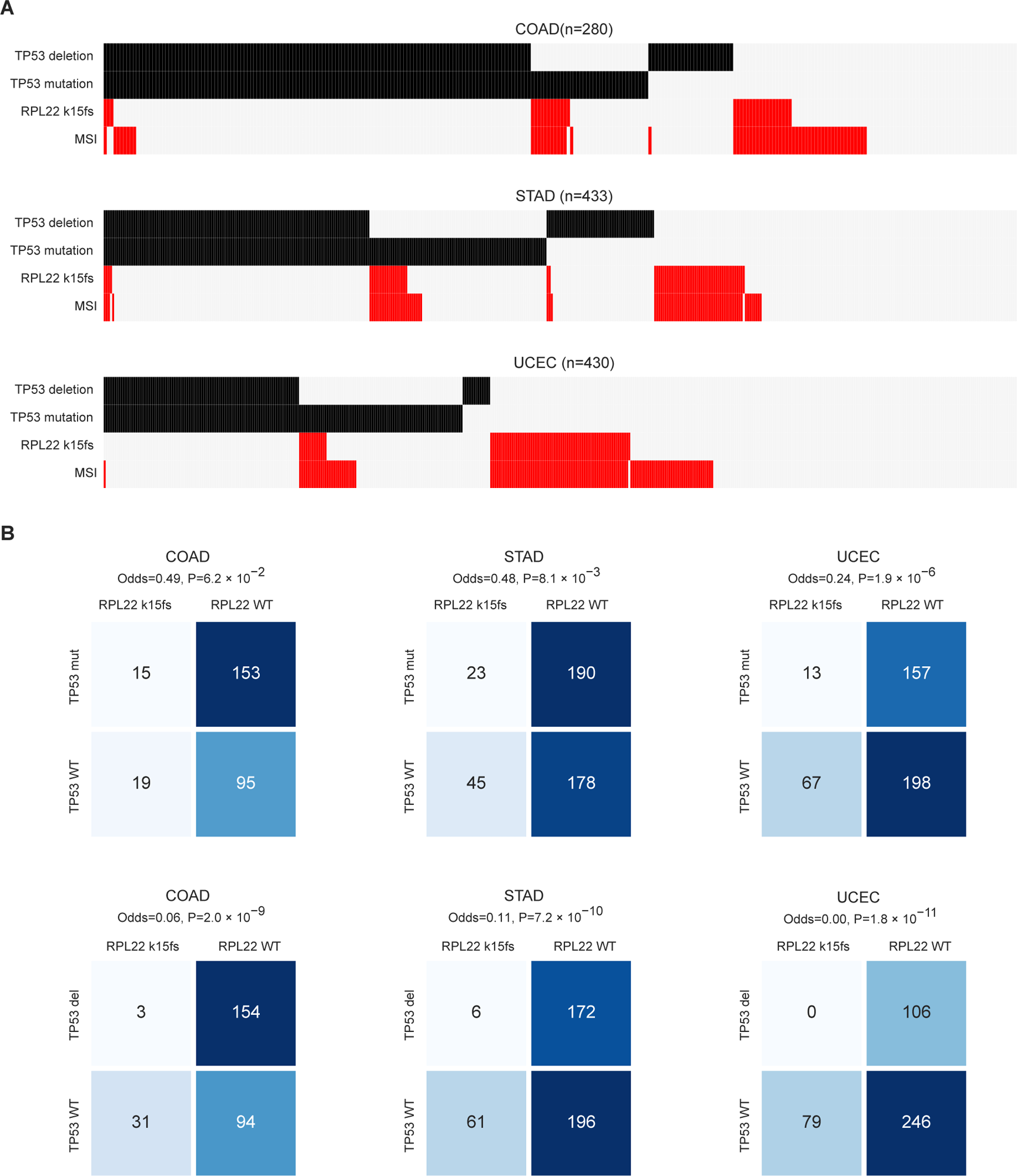
*RPL22* frameshift mutations are mutually exclusive with deleterious *TP53* alterations. (**A**) Heatmap of TP53 deletions and mutations, *RPL22 k15fs* mutations, and MSI occurrences in COAD, STAD, and UCEC samples of TCGA. (**B**) Contingency tables between *TP53* mutation (top row) and copy number deletion (bottom row) and *RPL22 k15fs* mutations in COAD (first column), STAD (second column), and UCEC (third column) samples of TCGA. Odds ratio indicates the relative frequency of *RPL22 k15fs* mutations between *TP53*-altered samples. P-value computed with Fisher’s exact test.

**Figure S2.**
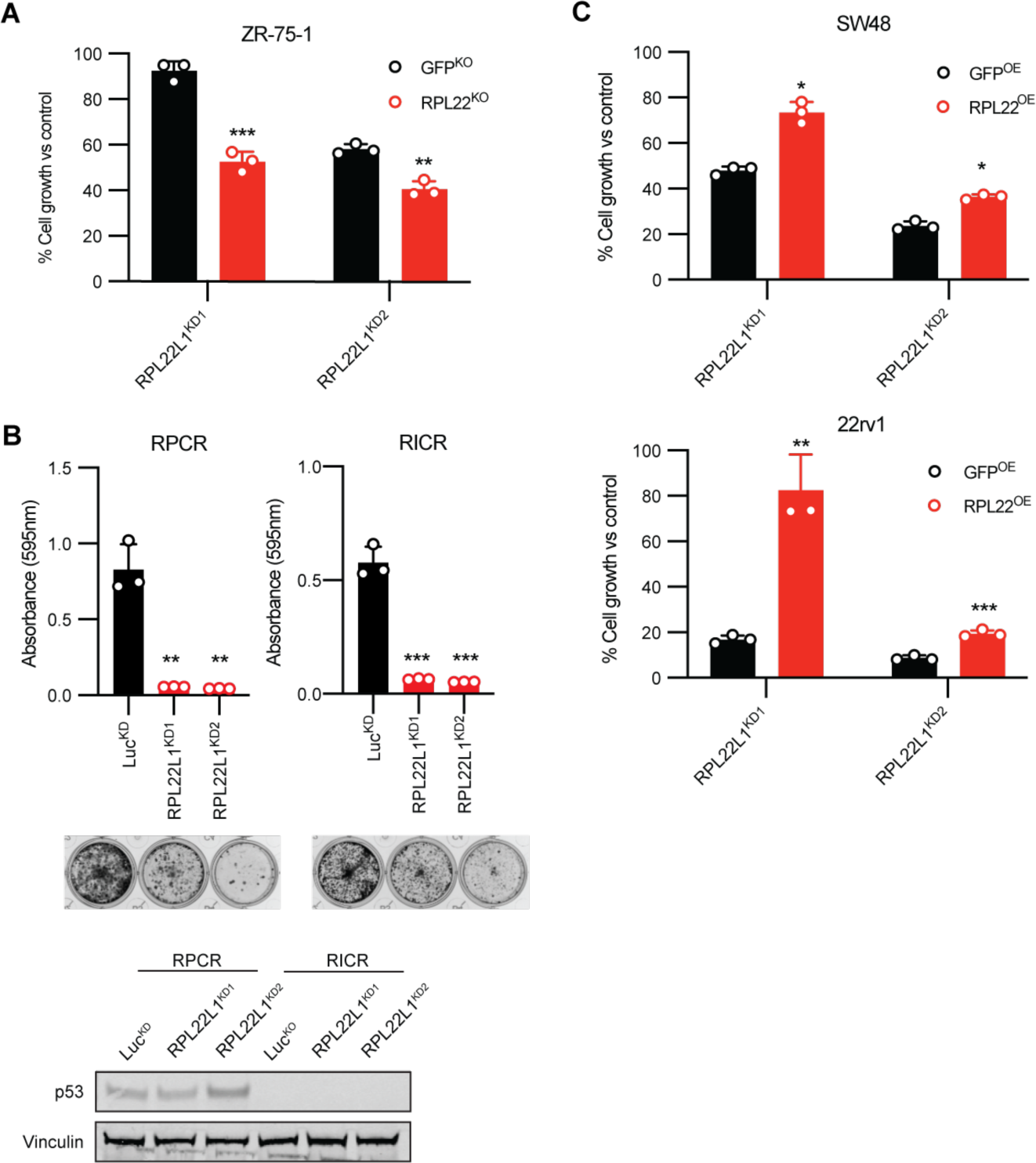
*RPL22L1* loss is rescued by *RPL22* expression. (**A**) Focus formation assay of cell lines (ZR-75-1, 7.5k cells/well) with CRISPR-cas9 knockout of *RPL22* and shRNA knockdown of *RPL22L1*. Crystal violet staining was quantitated 10-14 days post-seeding and compared to control by Student t test (p-value: ZR-75-1 RPL22^KO^ RPL22L1^KD1^ = 0.0003; ZR-75-1 RPL22^KO^ RPL22L1^KD2^ = 0.0019). Error bars represent the mean ± S.D of three replicates (* p < 0.05, ** p < 0.01, *** p < 0.001, **** p < 0.0001.) Data shown are representative figures of two independent experiments. (**B**) Focus formation assay and Western blot of *RPL22* mutant cell lines with wildtype P53 (RPCR) or mutant p53 (RICR) and subsequent shRNA knockdown of *RPL22L1*. Crystal violet staining was quantitated 7 days post-seeding and compared to control by Student t test (p-value: RPCR RPL22L1^KD1^ = 0.0014; RPCR RPL22L1^KD2^ = 0.0013; RICR RPL22L1^KD1^ = 0.0002; RICR RPL22L1^KD2^ = 0.0002). Error bars represent the mean ± S.D of three replicates (* p < 0.05, ** p < 0.01, *** p < 0.001, **** p < 0.0001.) Data shown are representative of three independent experiments. Blot immunostained for p53 (30ug protein/lane). (**C**) Focus formation assay of *RPL22* mutant cells with overexpression of *RPL22* and knockdown of *RPL22L1*. Crystal violet staining was quantitated 10-12 days post seeding and compared to control by Student t test (p-value: SW48 RPL22^OE^ RPL22L1^KD1^ = 0.0138; SW48 RPL22^OE^ RPL22L1^KD2^ = 0.0170; 22rv1 RPL22^OE^ RPL22L1^KD1^ = 0.00196; 22rv1 RPL22^OE^ RPL22L1^KD2^ = 0.0007). Error bars represent the mean ± S.D of three replicates (* p < 0.05, ** p < 0.01, *** p < 0.001, **** p < 0.0001.) Data shown are representative of two independent experiments.

**Figure S3.**
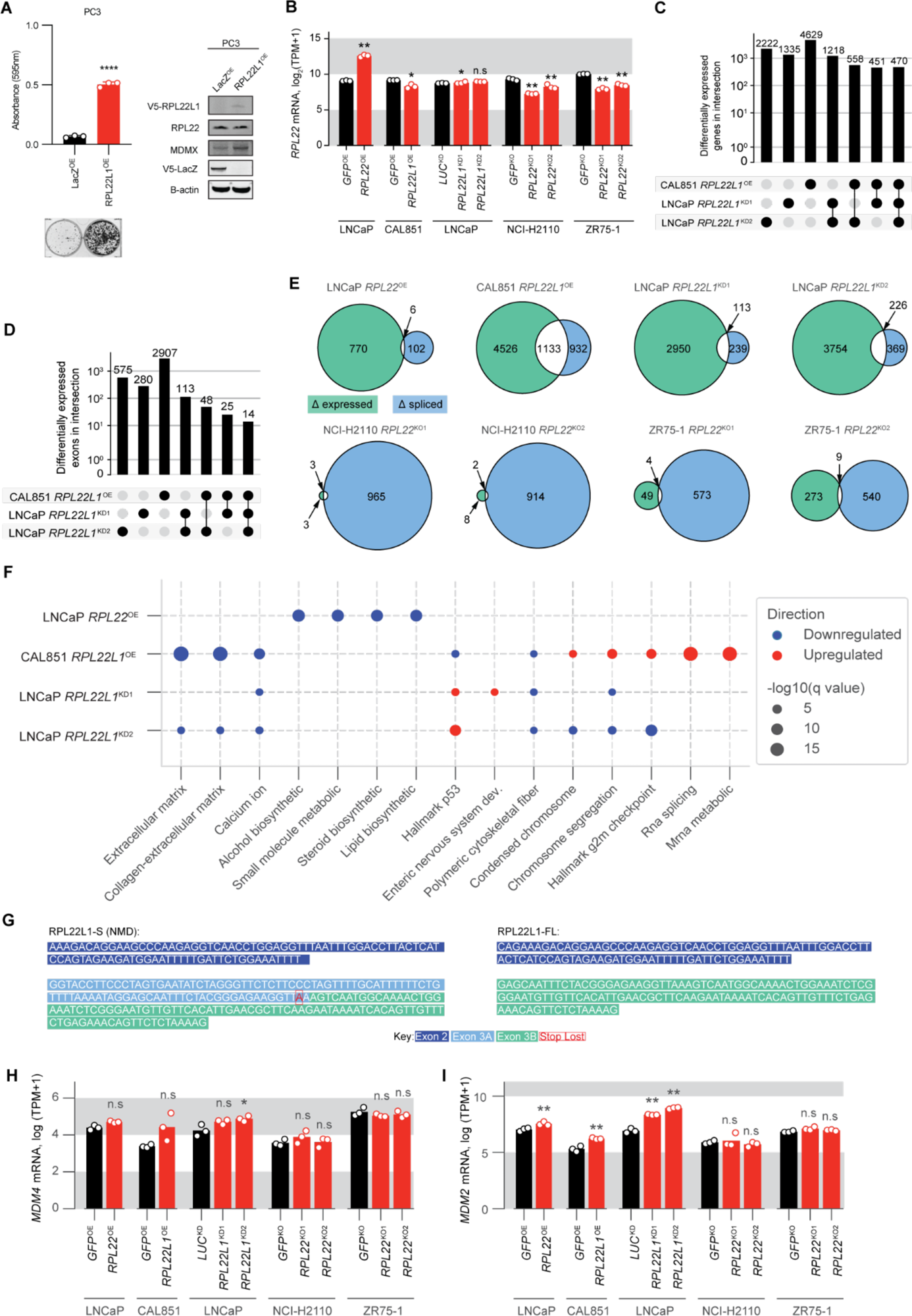
*RPL22L1* alters the expression of a broad array of genes and pathways. (**A**) Focus formation assay and Western blot of *RPL22* mutant cell lines with overexpression of *RPL22L1*. Crystal violet staining was quantitated 5 days post-seeding and compared to control by Student t test (p-value: PC3 RPL22L1^OE^ = <0.0001). Error bars represent the mean ± S.D of three replicates (* p < 0.05, ** p < 0.01, *** p < 0.001, **** p < 0.0001.) Data shown are representative of three independent experiments. Blot immunostained for V5-RPL22L1, RPL22, MDMX and V5-LacZ (30ug protein/lane). (**B**) Total *RPL22* mRNA expression across experiments. (**C**) Common differentially expressed genes across *RPL22L1* knockdown experiments. (**D**) Common splicing changes across *RPL22L1* overexpression and knockdown experiments. (**E**) Overlaps between differentially expressed and spliced genes across *RPL22* and *RPL22L1* modulation experiments. (**F**) Top enriched gene sets across experiments. (**G**) Cryptic splice site in *RPL22L1* exon 3. Sequence extracted from Ensembl. (**H**) Total *MDM4* mRNA expression across experiments. (**I**) Total *MDM2* mRNA expression across experiments.

**Figure S4.**
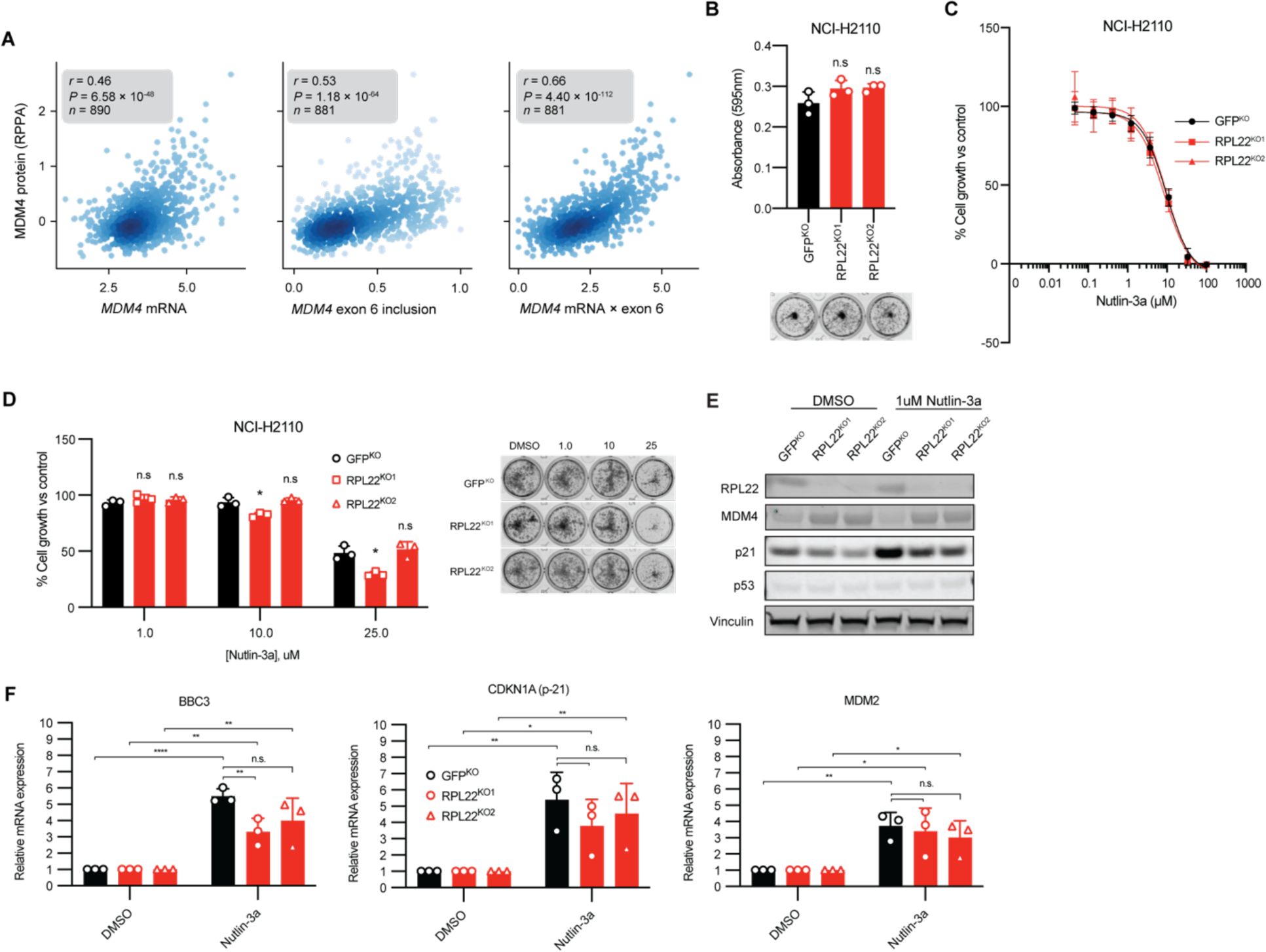
Sensitivity to Nutlin-3a is dependent on p53 status. (**A**) Predictive power of *MDM4* mRNA levels, exon 6 inclusion levels, and their product versus MDM4 protein levels. (**B**) Focus formation assay of cell line (NCI-H2110, 7.5k cells/well) with CRISPR-Cas9 knockout of *RPL22*. Crystal violet staining was quantitated 10-14 days post seeding and compared to control by Student t test (p-value: NCI-H2110 RPL22^KO1^ = 0.1510; NCI-H2110 RPL22^KO2^ = 0.0915). Error bars represent the mean ± S.D of three replicates (* p < 0.05, ** p < 0.01, *** p < 0.001, **** p < 0.0001.) Data shown are representative of at least three independent experiments. (**C**) Nutlin-3a treatment of cell lines (NCI-H2110, 2000 cells/well) with CRISPR-cas9 knockout of *RPL22*. Luminescence was quantitated after 6 days post-seeding and compared to control by Student t-test. Error bars represent the mean ± S.D of six replicates. Data shown are representative of three independent experiments. (**D**) Focus formation assay with nutlin-3a treatment of cell line (NCI-H2110, 7.5k cells/well) with CRISPR-cas9 knockout of *RPL22*. Crystal violet staining was quantitated 14 days post-seeding and compared to control by Student t test (p-value: 1.0 uM nutlin-3a: NCI-H2110 RPL22^KO1^ = 0.2484; NCI-H2110 RPL22^KO2^ = 0.2762; 10.0 uM nutlin-3a: NCI-H2110 RPL22^KO1^ = 0.0149; NCI-H2110 RPL22^KO2^ = 0.4790; 25.0 uM nutlin-3a: NCI-H2110 RPL22^KO1^ = 0.0079; NCI-H2110 RPL22^KO2^ = 0.5992). Error bars represent the mean ± S.D of three replicates (* p < 0.05, ** p < 0.01, *** p < 0.001, **** p < 0.0001.) Data shown are representative figures of three independent experiments. (**E**) Western blot of cell line (ZR-75-1) with CRISPR-Cas9 knockout of *RPL22* with immunostaining for p53 and p21 after 1 week of 1uM nutlin-3a treatment (25ug protein/lane). (**F**) mRNA expression of downstream targets of p53 in MSS ZR-75-1 cells with knockdown of *RPL22* and 1 week treatment with Nutlin-3a. Error bars represent the mean ± S.D of three replicates (* p < 0.05, ** p < 0.01, *** p < 0.001, **** p < 0.0001.) Data shown are representative figures of three independent experiments.

**Figure S5.**
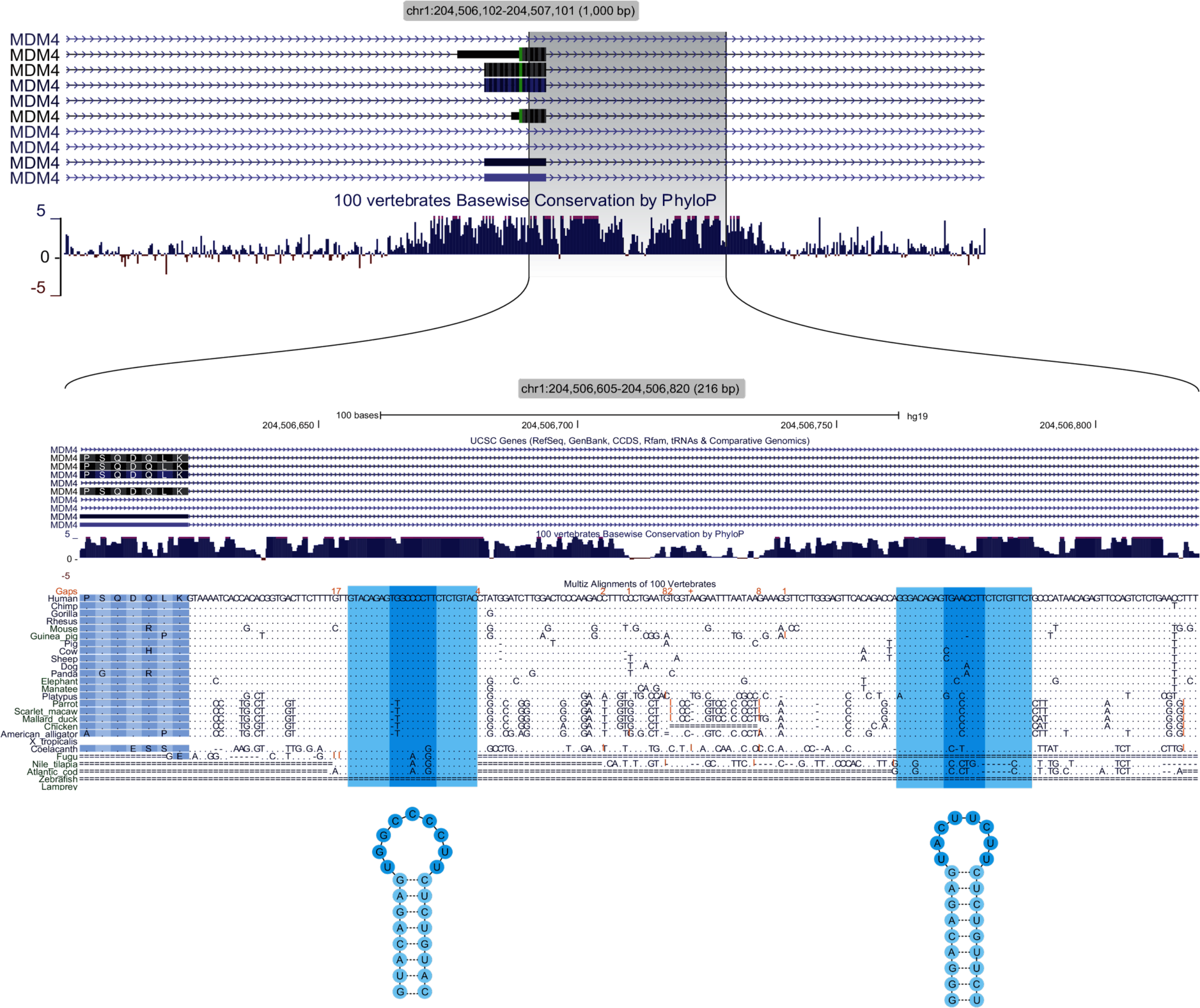
Predicted hairpin structures in the intronic regions of *MDM4* proximal to exon 6.

**Figure S6.**
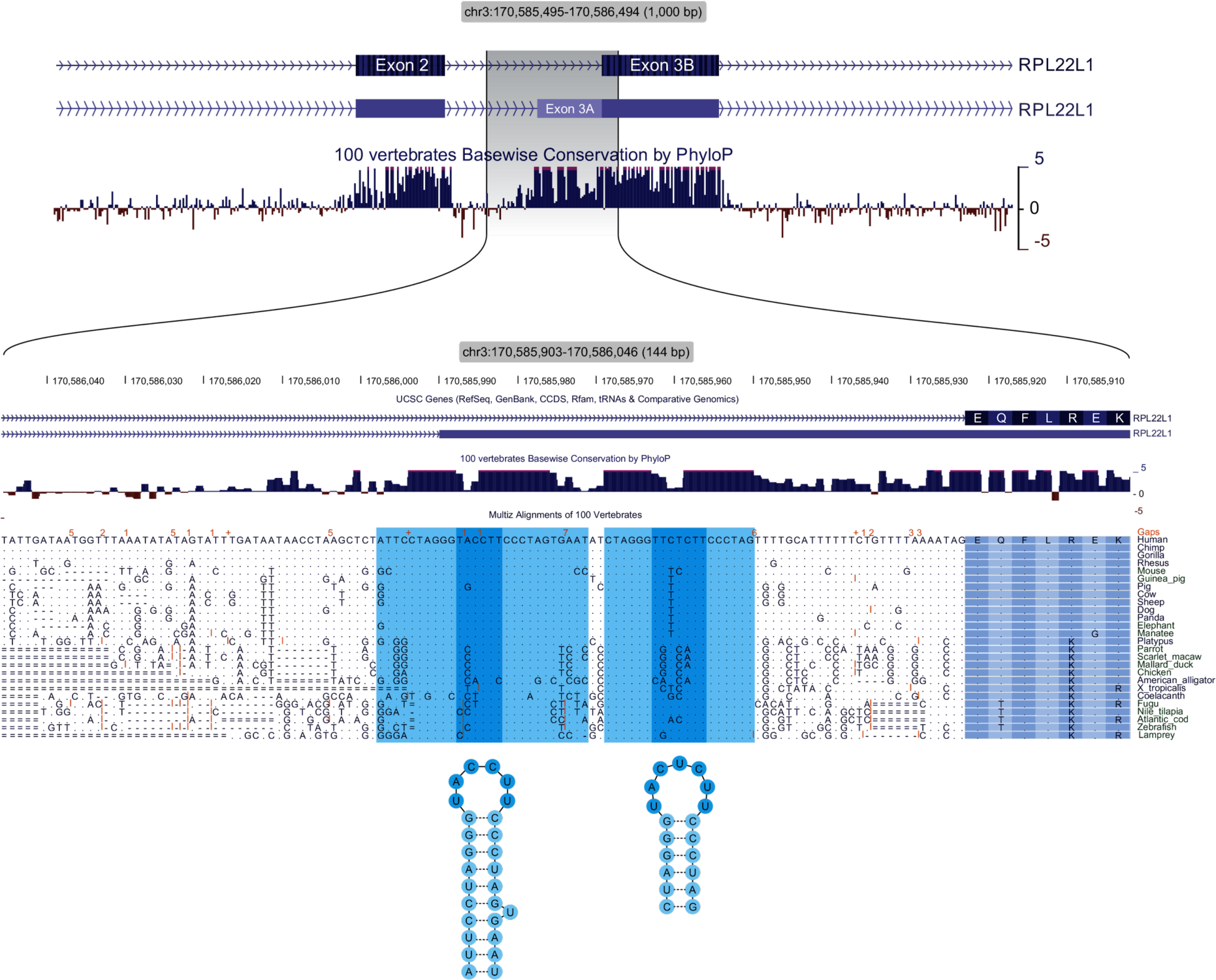
Predicted hairpin structures in the intronic regions of *RPL22L1* proximal to exon 3a.

**Figure S7.**
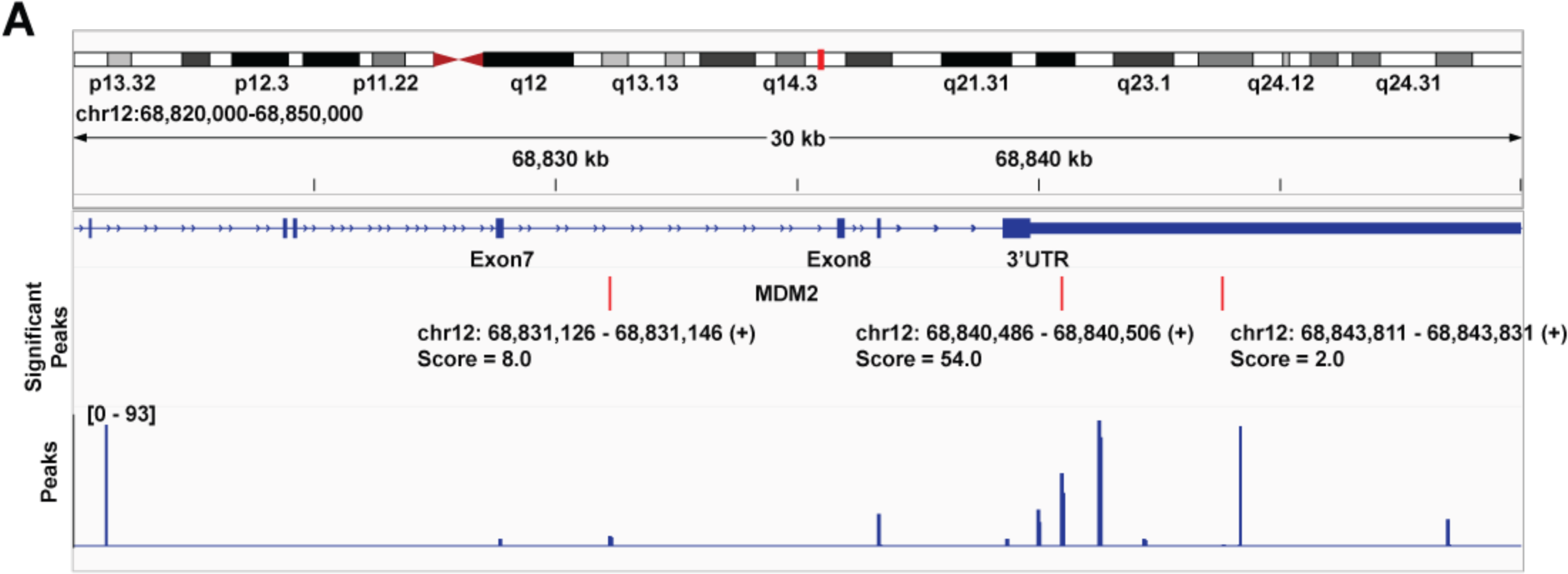
Characterization of RPL22 binding preference to MDM2 with CLIP-seq. (**A**) Binding of RPL22 to *MDM2* as determined by peak calling on CLIP-seq measurements from ZR-75-1 cells.

**Table S1.** Cancer cell lines used in this study.

| **Table S1. Cancer cell lines used in this study** |  |  |  |  |  |
| --- | --- | --- | --- | --- | --- |
| Cell line | Tissue | MSI status | *TP53* status | *RPL22* status | Source of RNA-seq data |
| SW48 | Colon | + | wt | mt | 41 |
| LNCaP | Prostate | + | wt | mt | 42 |
| 22rv1 | Prostate | + | wt | mt | 42 |
| SKMEL2 | Skin | + | mt | mt | 5 |
| RPCR | Colon | + | wt | mt | 41 |
| RICR | Colon | + | mt | mt | 41 |
| CAL851 | Breast | – | mt | wt | 43 |
| NCI-H2110 | Lung | – | mt | wt | 43 |
| ZR75-1 | Breast | – | wt | wt | 43 |
| MCF7 | Breast | – | wt | wt | 44 |
| C32 | Skin | – | wt | wt | 43 |
| PC3 | Prostate | – | mt | wt | 42,43 |
| Key: + = MSI; – = MSS; wt = wildtype; mt = mutated |  |  |  |  |  |

**Table S2.**
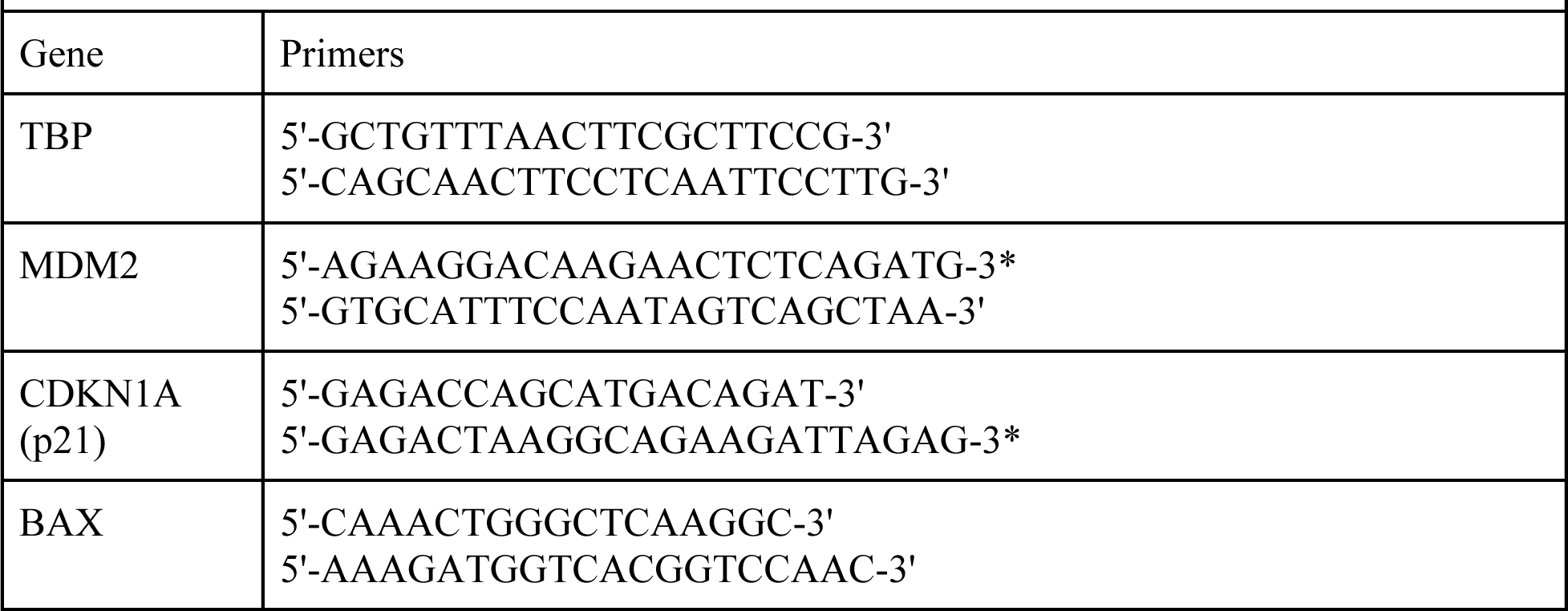
Primers used for qPCR.

**Data S1: MSI-H and *RPL22fs* mutation frequency in CCLE and TCGA data.** Provided as supplementary excel file.

**Data S2:**
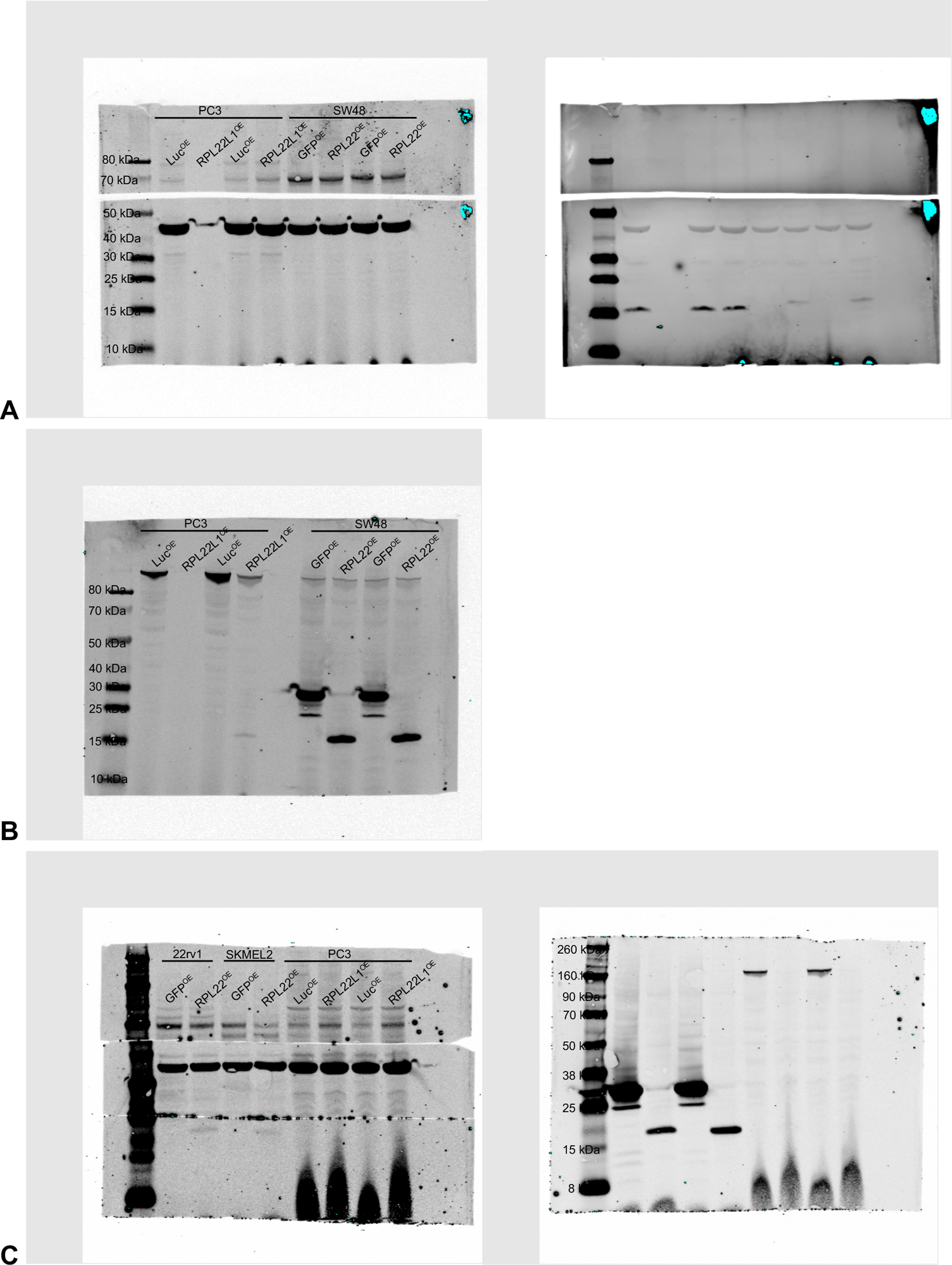
Figure 1i and S3a full western blot of cell lines with either *RPL22* or *RPL22L1* overexpression with immunostaining for RPL22, MDM4, and b-actin. (**A**) Multiple replicates of PC3 and SW48 lysates were loaded with 30ug protein per lane. The membrane was probed with the following primary antibodies: MDMX (Bethyl, A300-287A-M; top left), b-Actin (13E5) (4970S, Cell Signaling; bottom left) and RPL22-52 (sc-136413, Santa Cruz; right bottom). On the left is the Thermo Scientific Spectra Multicolor Broad Range Protein Ladder (26634). (**B**) The same PC3 and SW48 lysates were loaded on a subsequent day with 30ug protein per lane. The membrane was probed with the following primary antibodies: Vinculin (V9131, Sigma) and V5 to detect V5 tagged GFP, RPL22 and RPL22L1 (R96025, Fisher Scientific). On the left is the Thermo Scientific Spectra Multicolor Broad Range Protein Ladder (26634). (**C**) 22rv1, SKMEL2 and PC3 lysates were loaded with 30ug protein per lane. The membrane was probed with the following primary antibodies: MDMX (Bethyl, A300-287A-M; top left), b-Actin (13E5) (4970S, Cell Signaling; middle left), RPL22-52 (sc-136413, Santa Cruz; left bottom) and V5 to detect V5 tagged GFP, lacZ and RPL22. (R96025, Fisher Scientific; right). On the left is the Chameleon® Duo Pre-stained Protein Ladder (928-60000).

**Data S3.**
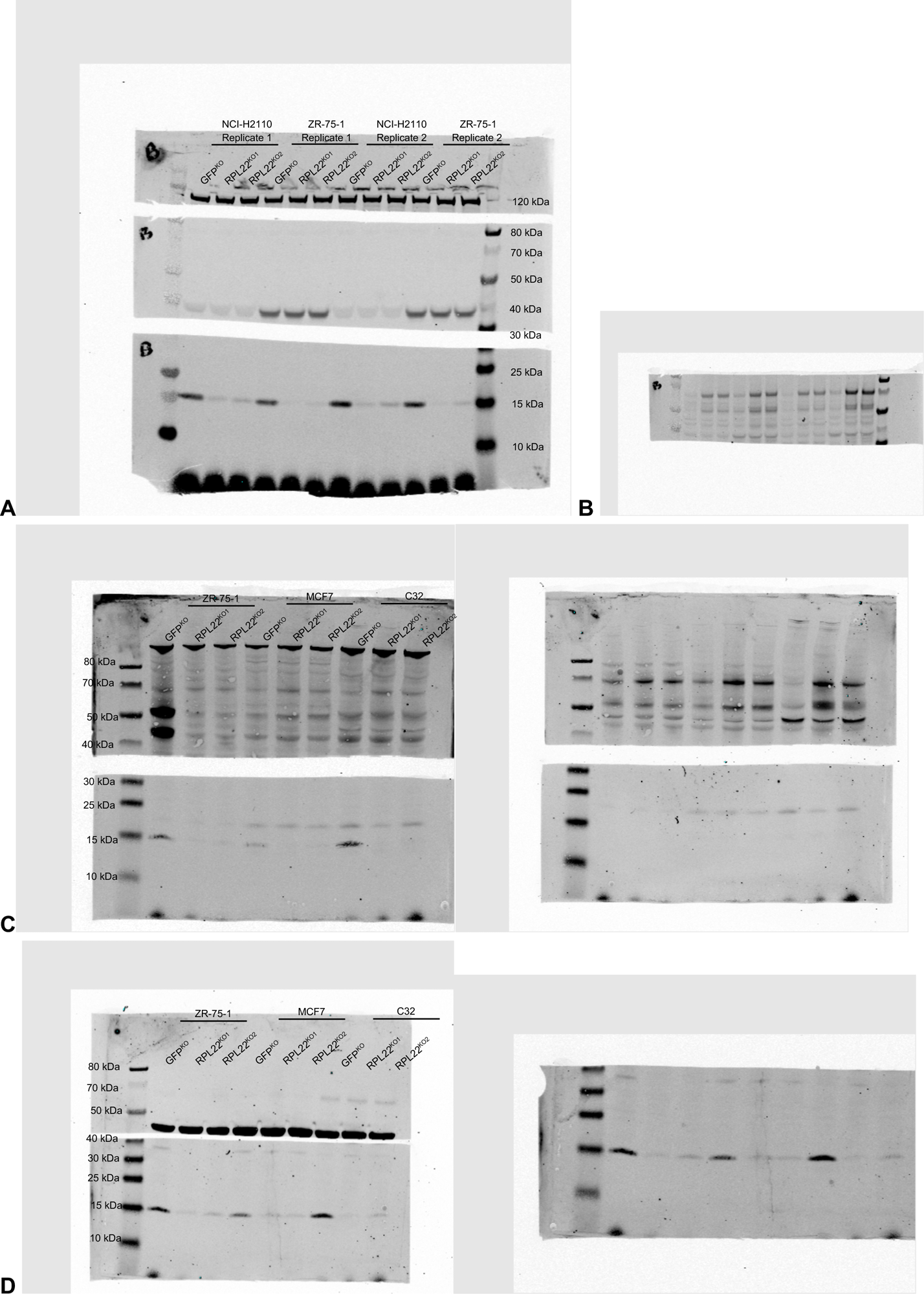
Figure 3a full western blot of cell lines with CRISPR-Cas9 knockout of *RPL22* with immunostaining for MDM4. (**A**) Two biological replicates of NCI-H2110 and ZR-75-1 lysates were loaded with 30ug protein per lane. Membrane was first probed with the following primary antibodies: Vinculin (V9131, Sigma; top), MDMX (G-10) (sc-74467, Santa Cruz; middle), and RPL22-52 (sc-136413, Santa Cruz; bottom). On the right is the Thermo Scientific Spectra Multicolor Broad Range Protein Ladder (26634). (**B**) Middle section of the membrane was re-probed with MDMX (Bethyl, A300-287A-M). (**C**) ZR-75-1, MCF7 and C32 lysates were loaded with 30ug protein per lane. Membrane was probed with the following primary antibodies: Vinculin (V9131, Sigma; top left), MDMX (G-10) (sc-74467, Santa Cruz; top right), p21 Waf1/Cip1 (DCS60) (2947S, Cell Signaling; bottom right) and RPL22-52 (sc-136413, Santa Cruz; bottom left). On the left is the Thermo Scientific Spectra Multicolor Broad Range Protein Ladder (26634). (**D**) The same ZR-75-1, MCF7 and C32 lysates were loaded again with 30ug protein per lane. Membrane was probed with the following primary antibodies: b-Actin (13E5) (4970S, Cell Signaling; top left), p53 (DO-1) (sc-126, Santa Cruz; bottom), and RPL22-52 (sc-136413, Santa Cruz; bottom). The bottom section of the gel was imaged twice (right). On the left is the Thermo Scientific Spectra Multicolor Broad Range Protein Ladder (26634).

**Data S4.**
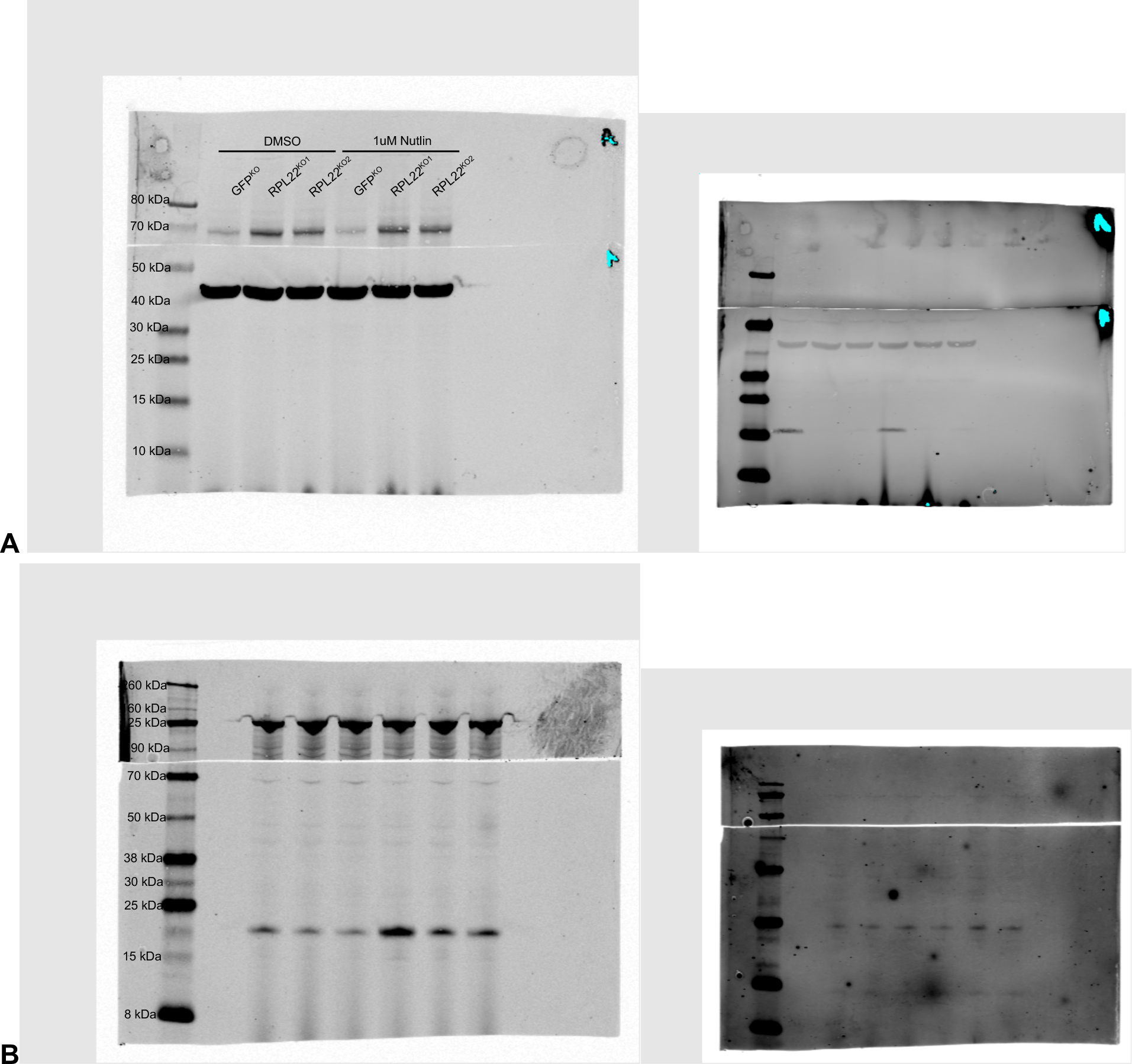
Figure 3f full western blot of cell line with CRISPR-Cas9 knockout of *RPL22* with immunostaining for p53 and p21 after 1 week of 1uM nutlin-3a treatment. (**A**) ZR-75-1 lysates were loaded with 30ug protein per lane. The membrane was probed with the following primary antibodies: MDMX (Bethyl, A300-287A-M; top left), b-Actin (13E5) (4970S, Cell Signaling; bottom left) and RPL22-52 (sc-136413, Santa Cruz; right bottom). On the left is the Thermo Scientific Spectra Multicolor Broad Range Protein Ladder (26634). (**B**) The same ZR-75-1 lysates were loaded with 30ug protein per lane on a subsequent day. The membrane was probed with the following primary antibodies: Vinculin (V9131, Sigma; top left), p21 Waf1/Cip1 (DCS60) (2947S, Cell Signaling; bottom left) and p53 (DO-1) (sc-126, Santa Cruz; bottom right). On the left is the Chameleon® Duo Pre-stained Protein Ladder (928-60000).

**Data S5.**
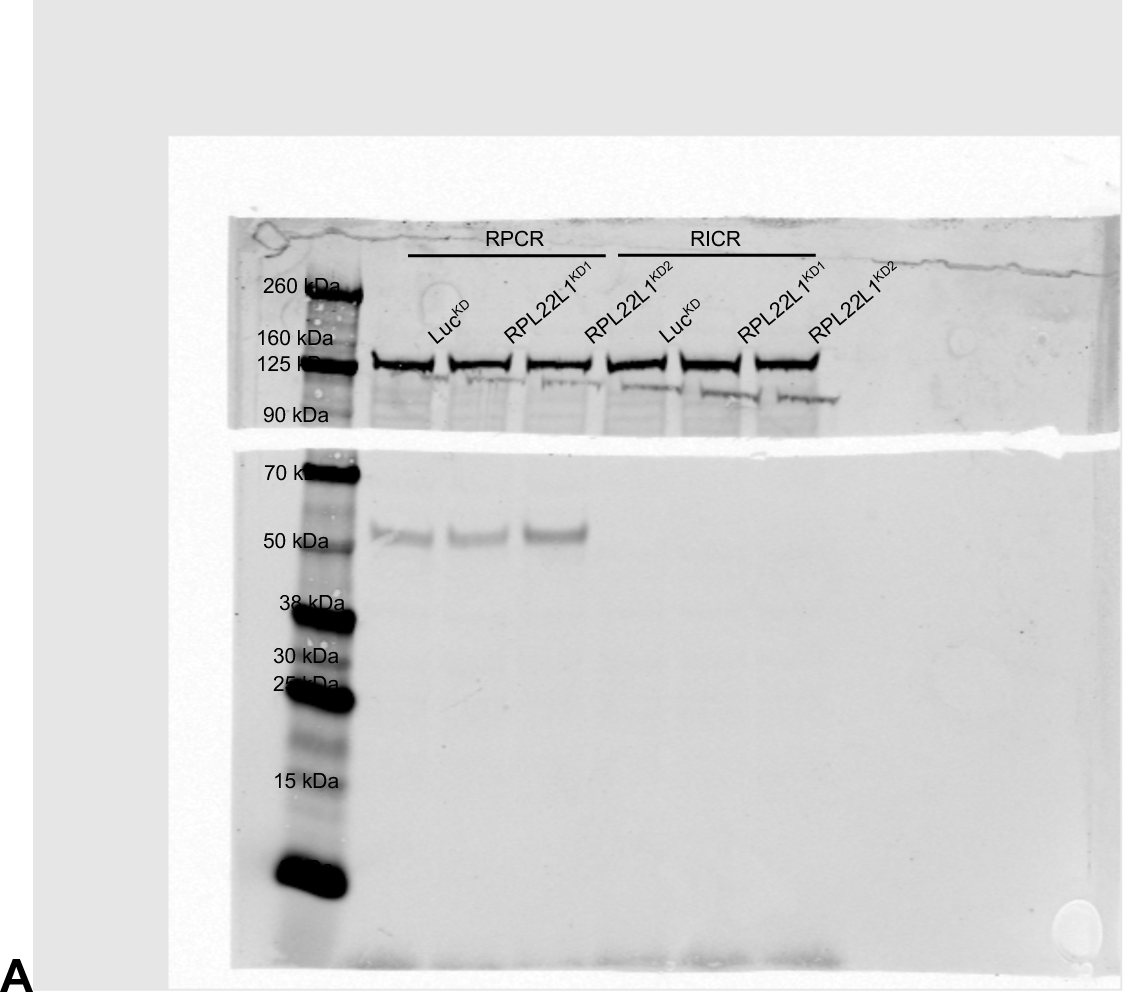
Figure S2c full western blot of RPL22 mutant cell lines with wildtype p53 (RPCR) or mutant p53 (RICR) with immunostaining for TP53. (**A**) Lysates were loaded with 30ug protein per lane. The membrane was probed with Vinculin (V9131, Sigma; top) and p53 (DO-1) (sc-126, Santa Cruz; bottom). On the left is the Chameleon® Duo Pre-stained Protein Ladder (928-60000).

**Data S6:**
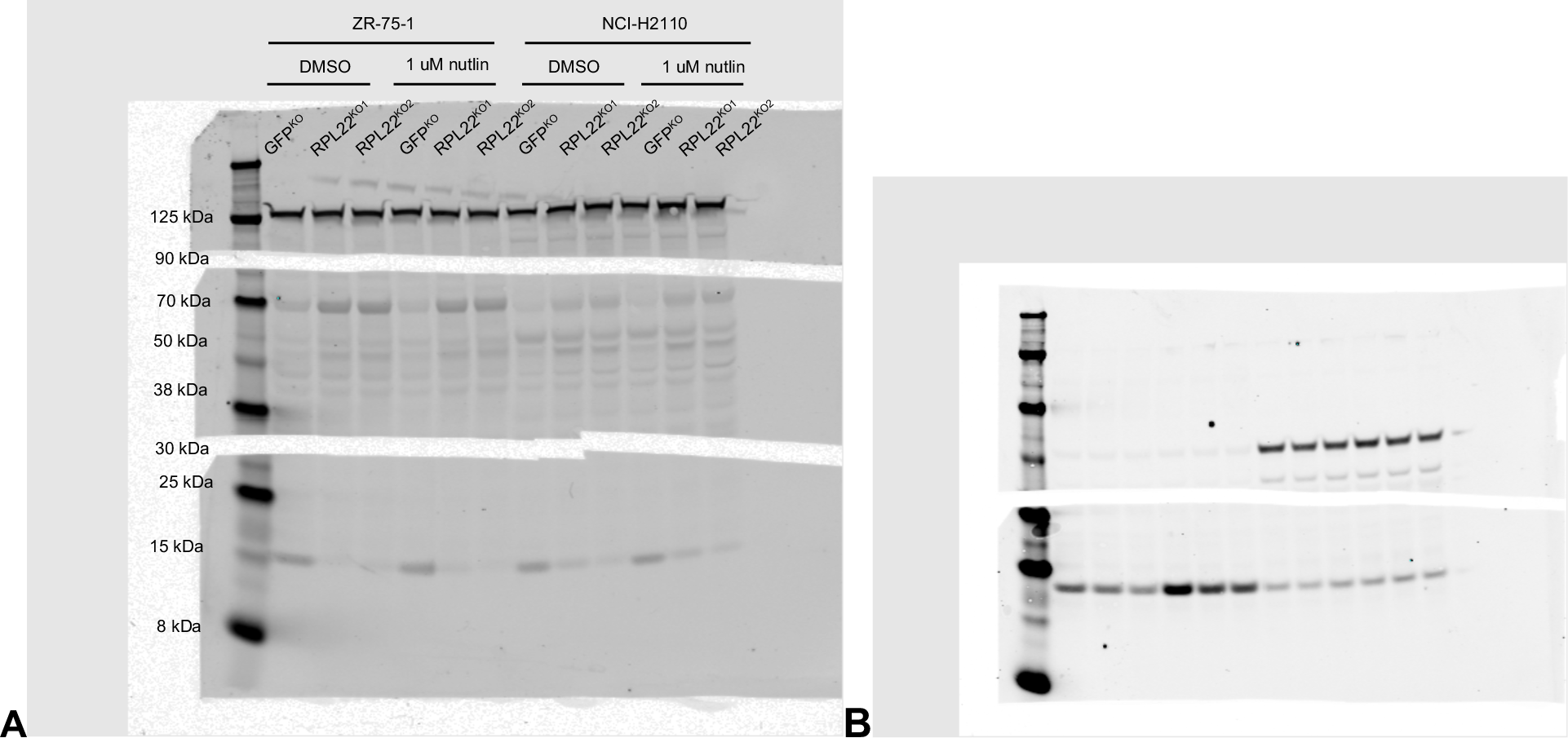
Figure S4d full western blot of cell lines with CRISPR-Cas9 knockout of *RPL22* with immunostaining for p53 and p21 after 1 week of 1uM nutlin-3a treatment. (**A**) ZR-75-1 and NCI-H2110 lysates were loaded with 25ug protein per lane. The membranes were probed with the following primary antibodies: Vinculin (V9131, Sigma; top), MDMX (Bethyl, A300-287A-M; middle), and RPL22-52 (sc-136413, Santa Cruz; bottom). On the left is the Chameleon® Duo Pre-stained Protein Ladder (928-60000). (**B**) The same lysates were loaded with 25ug protein per lane on the same day. The membrane was probed with the following primary antibodies: p53 (DO-1) (sc-126, Santa Cruz; top) and p21 Waf1/Cip1 (DCS60) (2947S, Cell Signaling; bottom). On the left is the Chameleon® Duo Pre-stained Protein Ladder (928-60000).

## References

1. Maruvka, Y. E. et al. Analysis of somatic microsatellite indels identifies driver events in human tumors. Nat. Biotechnol. 35, 951–959 (2017).

2. Eso, Y., Shimizu, T., Takeda, H., Takai, A. & Marusawa, H. Microsatellite instability and immune checkpoint inhibitors: toward precision medicine against gastrointestinal and hepatobiliary cancers. J. Gastroenterol. 55, 15–26 (2020).

3. Cortes-Ciriano, I., Lee, S., Park, W.-Y., Kim, T.-M. & Park, P. J. A molecular portrait of microsatellite instability across multiple cancers. Nat. Commun. 8, 15180 (2017).

4. Baugh, E. H., Ke, H., Levine, A. J., Bonneau, R. A. & Chan, C. S. Why are there hotspot mutations in the TP53 gene in human cancers? Cell Death Differ. 25, 154–160 (2018).

5. Ghandi, M. et al. Next-generation characterization of the Cancer Cell Line Encyclopedia. Nature 569, 503–508 (2019).

6. Novetsky, A. P. et al. Frequent mutations in the RPL22 gene and its clinical and functional implications. Gynecol. Oncol. 128, 470–474 (2013).

7. Ryan, M. et al. TCGASpliceSeq a compendium of alternative mRNA splicing in cancer. Nucleic Acids Res. 44, D1018–D1022 (2016).

8. Zhang, Y. et al. Ribosomal Proteins Rpl22 and Rpl22l1 Control Morphogenesis by Regulating Pre-mRNA Splicing. Cell Rep. 18, 545–556 (2017).

9. Fahl, S. P., Harris, B., Coffey, F. & Wiest, D. L. Rpl22 Loss Impairs the Development of B Lymphocytes by Activating a p53-Dependent Checkpoint. J. Immunol. 194, 200–209 (2015).

10. Rao, S. et al. Ribosomal Protein Rpl22 Controls the Dissemination of T-cell Lymphoma. Cancer Res. 76, 3387–3396 (2016).

11. Rao, S. et al. RPL22L1 induction in colorectal cancer is associated with poor prognosis and 5-FU resistance. PLOS ONE 14, e0222392 (2019).

12. Wu, N. et al. Ribosomal L22-like1 (RPL22L1) Promotes Ovarian Cancer Metastasis by Inducing Epithelial-to-Mesenchymal Transition. PLOS ONE 10, e0143659 (2015).

13. Del Toro, N. et al. Ribosomal protein RPL22/eL22 regulates the cell cycle by acting as an inhibitor of the CDK4-cyclin D complex. Cell Cycle 18, 759–770 (2019).

14. O’Leary, M. N. et al. The Ribosomal Protein Rpl22 Controls Ribosome Composition by Directly Repressing Expression of Its Own Paralog, Rpl22l1. PLoS Genet. 9, e1003708 (2013).

15. Dewaele, M. et al. Antisense oligonucleotide–mediated MDM4 exon 6 skipping impairs tumor growth. J. Clin. Invest. 126, 68–84 (2016).

16. Rallapalli, R., Strachan, G., Cho, B., Mercer, W. E. & Hall, D. J. A Novel MDMX Transcript Expressed in a Variety of Transformed Cell Lines Encodes a Truncated Protein with Potent p53 Repressive Activity. J. Biol. Chem. 274, 8299–8308 (1999).

17. Bezzi, M. et al. Regulation of constitutive and alternative splicing by PRMT5 reveals a role for Mdm4 pre-mRNA in sensing defects in the spliceosomal machinery. Genes Dev. 27, 1903–1916 (2013).

18. Bieging-Rolett, K. T. et al. Zmat3 Is a Key Splicing Regulator in the p53 Tumor Suppression Program. Molecular Cell 80, 452–469.e9 (2020).

19. McDonald, E. R. et al. Project DRIVE: A Compendium of Cancer Dependencies and Synthetic Lethal Relationships Uncovered by Large-Scale, Deep RNAi Screening. Cell 170, 577–592.e10 (2017).

20. Chan, E. M. et al. WRN helicase is a synthetic lethal target in microsatellite unstable cancers. Nature 568, 551–556 (2019).

21. Luo, E.-C. et al. Large-scale tethered function assays identify factors that regulate mRNA stability and translation. Nat. Struct. Mol. Biol. 27, 989–1000 (2020).

22. Vassilev, L. T. et al. In Vivo Activation of the p53 Pathway by Small-Molecule Antagonists of MDM2. Science 303, 844–848 (2004).

23. Cao, B. et al. Cancer-mutated ribosome protein L22 (RPL22/eL22) suppresses cancer cell survival by blocking p53-MDM2 circuit. Oncotarget 8, 90651–90661 (2017).

24. Haupt, S., Mejía-Hernández, J. O., Vijayakumaran, R., Keam, S. P. & Haupt, Y. The long and the short of it: the MDM4 tail so far. Journal of Molecular Cell Biology 11, 231–244 (2019).

25. Gabunilas, J. & Chanfreau, G. Splicing-Mediated Autoregulation Modulates Rpl22p Expression in Saccharomyces cerevisiae. PLOS Genet. 12, e1005999 (2016).

26. Larionova, T. D. et al. Alternative RNA splicing modulates ribosomal composition and determines the spatial phenotype of glioblastoma cells. Nat. Cell Biol. 24, 1541–1557 (2022).

27. Abrhámová, K. & Folk, P. Regulation of yeast RPL22B splicing depends on intact pre-mRNA context of the intron. http://biorxiv.org/lookup/doi/10.1101/814301 (2019) doi:10.1101/814301.

28. Petibon, C., Parenteau, J., Catala, M. & Elela, S. A. Introns regulate the production of ribosomal proteins by modulating splicing of duplicated ribosomal protein genes. Nucleic Acids Res. 44, 3878–3891 (2016).

29. Abrhámová, K. et al. Introns provide a platform for intergenic regulatory feedback of RPL22 paralogs in yeast. PLOS ONE 13, e0190685 (2018).

30. Sciarrillo, R. et al. Splicing modulation as novel therapeutic strategy against diffuse malignant peritoneal mesothelioma. EBioMedicine 39, 215–225 (2019).

31. Wang, E. et al. Targeting an RNA-Binding Protein Network in Acute Myeloid Leukemia. Cancer Cell 35, 369–384.e7 (2019).

32. Seiler, M. et al. Somatic Mutational Landscape of Splicing Factor Genes and Their Functional Consequences across 33 Cancer Types. Cell Rep. 23, 282–296.e4 (2018).

33. Kahles, A. et al. Comprehensive Analysis of Alternative Splicing Across Tumors from 8,705 Patients. Cancer Cell 34, 211–224.e6 (2018).

34. Georgilis, A. et al. PTBP1-Mediated Alternative Splicing Regulates the Inflammatory Secretome and the Pro-tumorigenic Effects of Senescent Cells. Cancer Cell 34, 85–102.e9 (2018).

35. Bonneville, R. et al. Landscape of Microsatellite Instability Across 39 Cancer Types. *JCO Precis*. Oncol. 1–15 (2017) doi:10.1200/po.17.00073.

36. Sanjana, N. E., Shalem, O. & Zhang, F. Improved vectors and genome-wide libraries for CRISPR screening. Nature Methods 11, 783–784 (2014)

37. Shalem, O. et al. Genome-Scale CRISPR-Cas9 Knockout Screening in Human Cells. Science 343, 84–87 (2014).

38. Shah, A., Qian, Y., Weyn-Vanhentenryck, S. M. & Zhang, C. CLIP Tool Kit (CTK): a flexible and robust pipeline to analyze CLIP sequencing data. Bioinformatics btw653 (2016) doi:10.1093/bioinformatics/btw653.

39. Robinson, J. T. et al. Integrative genomics viewer. Nat. Biotechnol. 29, 24–26 (2011).

40. Heinz, S. et al. Simple Combinations of Lineage-Determining Transcription Factors Prime cis-Regulatory Elements Required for Macrophage and B Cell Identities. Mol. Cell 38, 576– 589 (2010).

41. Ahmed, D. et al. Epigenetic and genetic features of 24 colon cancer cell lines. Oncogenesis 2, e71–e71 (2013).

42. Prensner, J. R. et al. Transcriptome sequencing across a prostate cancer cohort identifies PCAT-1, an unannotated lincRNA implicated in disease progression. Nat. Biotechnol. 29, 742–749 (2011).

43. DepMap. DepMap 19Q4 Public. figshare. Dataset. Broad (2019) doi:10.6084/m9.figshare.11384241.v2 doi:10.6084/M9.FIGSHARE.22765112.V2.

44. Edgren, H. et al. Identification of fusion genes in breast cancer by paired-end RNA-sequencing. Genome Biol. 12, R6 (2011).

